# Activity-based fluorescence screening uncovers host enzymatic responses to SARS-CoV-2 infection

**DOI:** 10.64898/2026.11.17.766338

**Authors:** Kyohhei Fujita, Yukiko Muramoto, Kazuya Shimura, Mako Kamiya, Hayato Wakabayashi, Miyuki Tani, Chiho Onishi, Yoshio Koyanagi, Takeshi Noda, Yasuteru Urano

## Abstract

Transcriptomic and proteomic studies have characterized host responses to SARS-CoV-2 infection, yet changes in endogenous enzymatic activities remain poorly understood. Here, we screened 89 enzyme-activatable fluorescence probes in living SARS-CoV-2-infected Calu-3 cells and identified 18 probes showing enhanced fluorescence, with some exhibiting multiplicity-of-infection-dependent responses. Neither SARS-CoV-2 M^pro^ nor PL^pro^ activated the tested probes, and several did not exhibit comparable fluorescence enhancements in influenza-virus-infected cells. Target identification and biochemical validation implicated calpain-1 (CAPN1) and puromycin-sensitive aminopeptidase (PSA) as major contributors to probe activation. A PSA-reactive fluorescence probe confirmed that SARS-CoV-2-infected living cells showed enhanced PSA activity, which was suppressed by puromycin. Immunofluorescence analysis revealed a strong positive correlation between PSA and SARS-CoV-2 spike protein signals. Notably, neither CAPN1 nor PSA was included among differentially expressed genes or proteins previously identified in SARS-CoV-2-infected cells. These findings highlight activity-based fluorescence screening as a complementary strategy to expression-based analyses for discovering infection-induced biomarker enzymes.

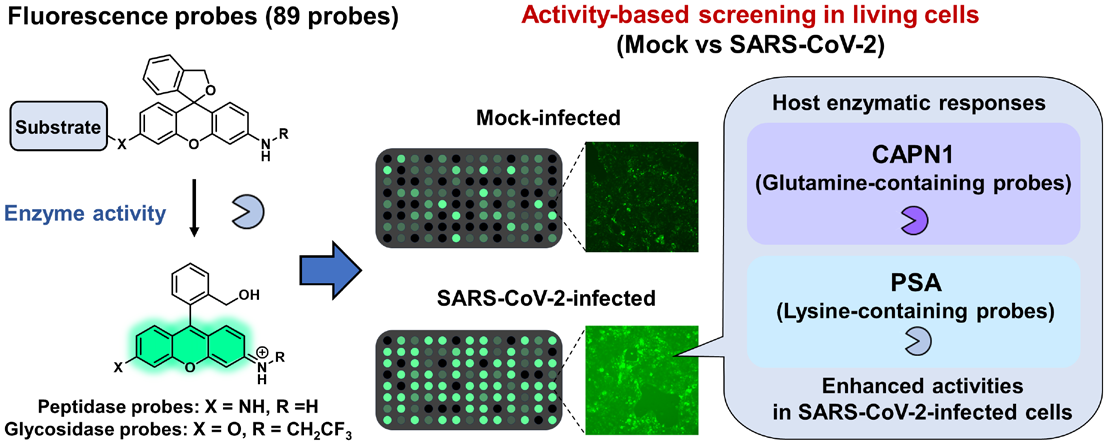

## Introduction

Severe acute respiratory syndrome coronavirus 2 (SARS-CoV-2), the causative agent of coronavirus disease 2019 (COVID-19),^1^ infects respiratory epithelial cells and induces extensive alterations in host-cell physiology.^2, 3^ Understanding these infection-induced changes is crucial for elucidating the molecular mechanisms of viral infection and may facilitate the identification of novel therapeutic targets and diagnostic biomarkers.^4-6^ Previous transcriptomic and proteomic studies have revealed widespread changes in host-cell processes, including metabolism, signaling pathways and stress responses, following SARS-CoV-2 infection.^6-9^ Despite these advances, however, infection-induced changes in endogenous enzymatic activities remain poorly characterized.

Proteolytic enzymes play essential roles in the SARS-CoV-2 life cycle. Viral proteases, including the main protease (M^pro^) and papain-like protease (PL^pro^), are responsible for processing viral polyproteins,^10, 11^ whereas host proteases, such as transmembrane serine protease 2 (TMPRSS2) and cathepsins, facilitate viral entry.^12^ Although the activities of individual host proteases have been investigated during SARS-CoV-2 infection,^13, 14^ systematic profiling of infection-induced changes in endogenous host-cell enzymatic activities across diverse enzyme classes, including aminopeptidases, proteases and glycosidases, remains limited. Importantly, because enzymatic activities are regulated by various factors, including post-translational modifications and the intracellular milieu, changes in gene or protein expression do not necessarily reflect enzyme activity.^15, 16^ Thus, approaches that directly measure enzymatic activity in living cells may provide complementary insights into functional host-cell alterations associated with viral infection.

Enzyme-activatable fluorescence probes offer a powerful approach for directly visualizing enzymatic activities in living cells.^17^ We have developed a library of approximately 400 fluorescence probes targeting peptidases and glycosidases with diverse peptide and sugar substrates.^18, 19^ By screening these probes in clinical cancer specimens, we have identified probe/biomarker enzyme pairs that enable cancer-selective fluorescence imaging across various cancer types.^20, 21^ Peptidase-reactive probes are designed based on hydroxymethyl rhodamine green (HMRG) conjugated to amino acid or dipeptide substrates (P1-HMRG or P2-P1-HMRG),^18, 22^ whereas glycosidase-reactive probes are constructed by introducing monosaccharide substrates into hydroxymethyl rhodol ethyltrifluoride (HMRef).^19, 23^ These probes exist in a non-fluorescent, intramolecularly spirocyclized form, but are converted to the highly fluorescent open form upon reaction with the target enzyme. Such fluorescence probe libraries provide an attractive platform for systematically profiling endogenous enzymatic activities in living cells.

Here, we applied activity-based fluorescence screening using a selected panel of 89 enzyme-activatable probes to SARS-CoV-2-infected Calu-3 cells, a well-established human respiratory epithelial cell model of SARS-CoV-2 infection, and identified multiple infection-induced increases in endogenous enzymatic activities. Subsequent target identification and biochemical validation implicated calpain-1 (CAPN1) and puromycin-sensitive aminopeptidase (PSA) as major enzymes contributing to distinct probe responses. Furthermore, we demonstrated fluorescence visualization of enhanced PSA activity in SARS-CoV-2-infected living cells. These findings highlight the potential of activity-based fluorescence screening to uncover functional host-cell alterations that may not be readily identifiable through conventional expression-based analyses.

## Results

### Identification of selectively activated probes in SARS-CoV-2-infected cells

To identify fluorescence probe/target enzyme pairs that exhibit enhanced fluorescence in SARS-CoV-2-infected cells, we first screened a panel of 89 fluorescence probes (77 peptidase-reactive probes and 12 glycosidase-reactive probes) in Calu-3 cells using live-cell fluorescence imaging (**Fig. 1a, Table S1**). These probes were selected from our fluorescence probe library.^18, 19^ For the peptidase-reactive probes, P1 residues were selected based on our previous observations of robust fluorescence activation in human cells,^18, 20^ whereas P2 residues were chosen to cover a range of physicochemical properties, including basic, acidic and neutral amino acids. We used Calu-3 cells as a well-established human respiratory epithelial model of SARS-CoV-2 infection because they endogenously express angiotensin-converting enzyme 2 (ACE2) and TMPRSS2, support viral replication, and exhibit infection-associated host-cell responses.^24, 25^ Calu-3 cells were infected with SARS-CoV-2 at a multiplicity of infection (MOI) of 1, and changes in fluorescence intensity were evaluated immediately (0 h) and 2 h after probe addition. We found that several probes showed a marked fluorescence increase specifically in SARS-CoV-2 infected Calu-3 cells, compared to mock-infected cells (**Fig. 1b, c, Fig. S1**). We defined primary hits as probes that exhibited both a high fluorescence intensity ratio between SARS-CoV-2-infected and mock-infected cells and a substantial fluorescence increase in infected cells. Specifically, probes with a fluorescence intensity ratio of SARS-CoV-2-infected to mock-infected cells (I/M ratio) of ≥1.2 at 2 h and a fluorescence increase in infected cells from 0 to 2 h (ΔFI) of ≥1.5 (a.u.) were classified as primary hits. Based on these criteria, 18 probes were identified as hits, including 16 peptidase-reactive probes and 2 glycosidase-reactive probes (**Fig. 1d**).

**Figure 1.**
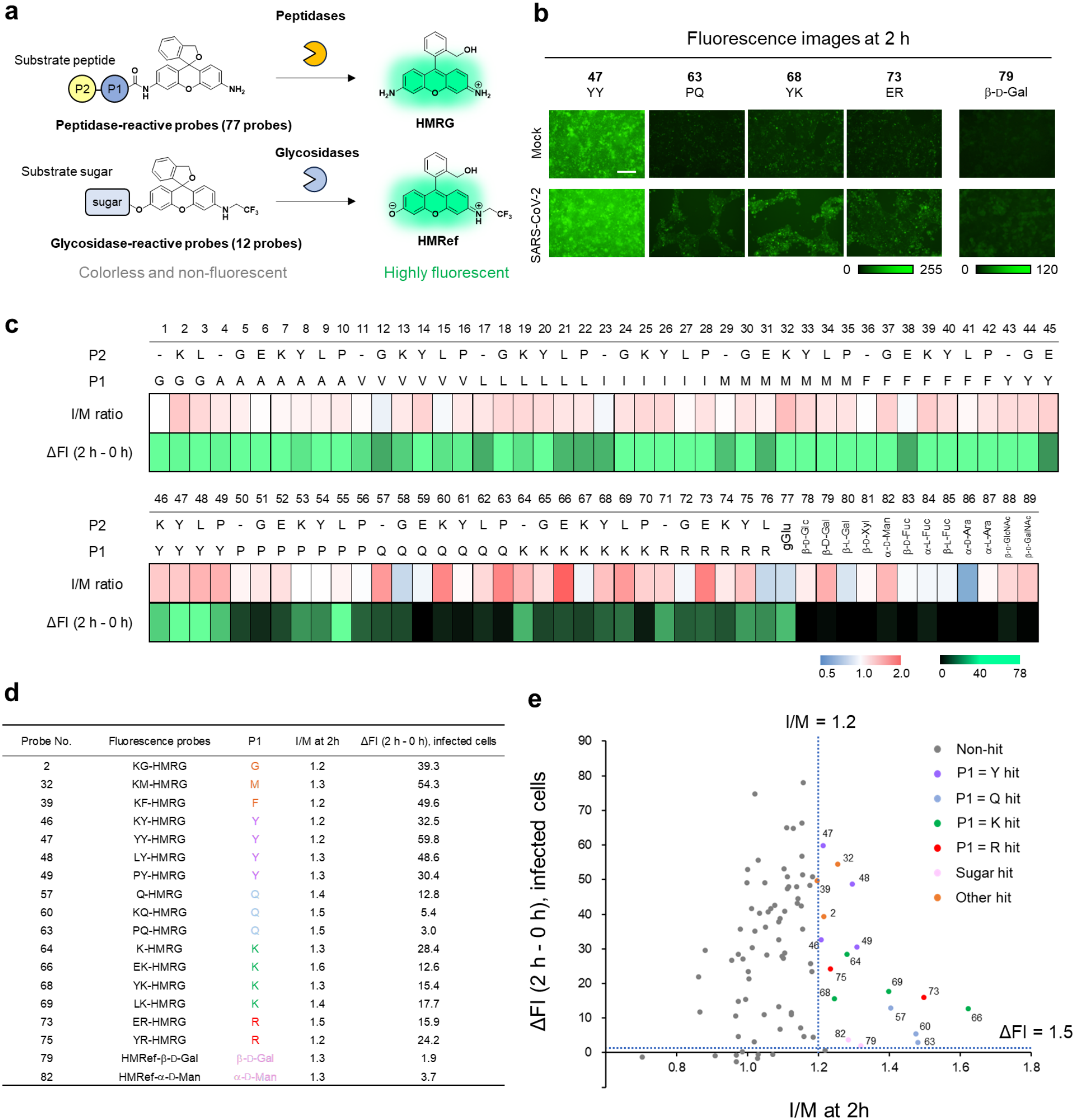
Activity-based screening of fluorescence probes in SARS-CoV-2-infected Calu-3 cells using live-cell imaging. (**a**) Reaction scheme of HMRG or HMRef-based activatable fluorescence probes targeting enzyme activities. A total of 89 probes were evaluated, of which 77 probes targeted aminopeptidases and proteases and 12 probes targeted glycosidases. (**b**) Representative fluorescence images of mock-infected and SARS-CoV-2-infected Calu-3 cells acquired 2 h after probe addition during activity-based screening. Calu-3 cells were infected with SARS-CoV-2 at an MOI of 1, and fluorescence probes were added at 24 h post-infection. Several probes exhibited higher fluorescence intensities in SARS-CoV-2-infected cells than in mock-infected cells. [fluorescence probe] = 10 μM. Scale bar, 200 μm. (**c**) Heat maps showing the fluorescence intensity ratios of SARS-CoV-2-infected cells to mock-infected cells (I/M ratios) at 2 h and the increases in fluorescence intensity from 0 to 2 h (ΔFI) in SARS-CoV-2-infected cells for all 89 probes. (**d**) List of 18 primary hit probes identified by the screening. (**e**) Scatter plot of the 89 evaluated probes showing the relationship between the I/M ratio at 2 h and ΔFI in SARS-CoV-2-infected cells. Primary hits were defined as probes with an I/M ratio ≥ 1.2 and ΔFI ≥ 1.5. Dashed lines indicate the thresholds for primary hit selection. Hit probes are color-coded according to their P1 amino acid residues or substrate classes.

Among the 16 peptidase-reactive probes, responsive substrates were observed across several P1 residues; however, probes bearing Q, K or R at the P1 position tended to show particularly pronounced differences between infected and mock-infected cells (**Fig. 1e**). Although several Y-containing probes also met the criteria, their I/M ratios were generally lower than those observed for the most responsive Q-, K-, and R-containing probes. These results suggested that substrates containing Q, K or R at P1 were particularly sensitive to the activity changes detected under the screening conditions. Furthermore, glycosidase-reactive probes such as HMRef-β-D-Gal and HMRef-α-D-Man also met the criteria and exhibited elevated fluorescence. None of these probes showed fluorescence activation in the presence of SARS-CoV-2 M^pro^ and PL^pro^ (**Fig. S2-3**), suggesting that the observed fluorescence enhancements reflect increased activity of endogenous host-cell enzymes associated with SARS-CoV-2 infection, rather than activation by viral protease. We therefore selected 9 representative probes, primarily those bearing Q, K or R at P1 and HMRef-β-D-Gal, based on their pronounced fluorescence responses and substrate diversity, to investigate whether their fluorescence responses are dependent on the MOI.

### Evaluation of MOI-dependent activation of fluorescence probes

To determine whether the observed fluorescence changes are MOI-dependent, Calu-3 cells were infected with SARS-CoV-2 at MOIs of 0, 0.1, 1, and 10, and live-cell fluorescence imaging was performed using 9 probes selected from the initial screening: Q-HMRG, KQ-HMRG, PQ-HMRG, K-HMRG, EK-HMRG, YK-HMRG, R-HMRG, ER-HMRG and HMRef-β-D-Gal. These probes showed greater fluorescence increases with increasing MOI, suggesting that their activation is positively correlated with SARS-CoV-2 infection (**Fig. 2a, b**). In particular, EK-HMRG, YK-HMRG and ER-HMRG exhibited approximately 2.5-fold higher fluorescence intensities in cells infected with SARS-CoV-2 at an MOI of 10 compared with uninfected cells (**Fig. 2c**).

**Figure 2.**
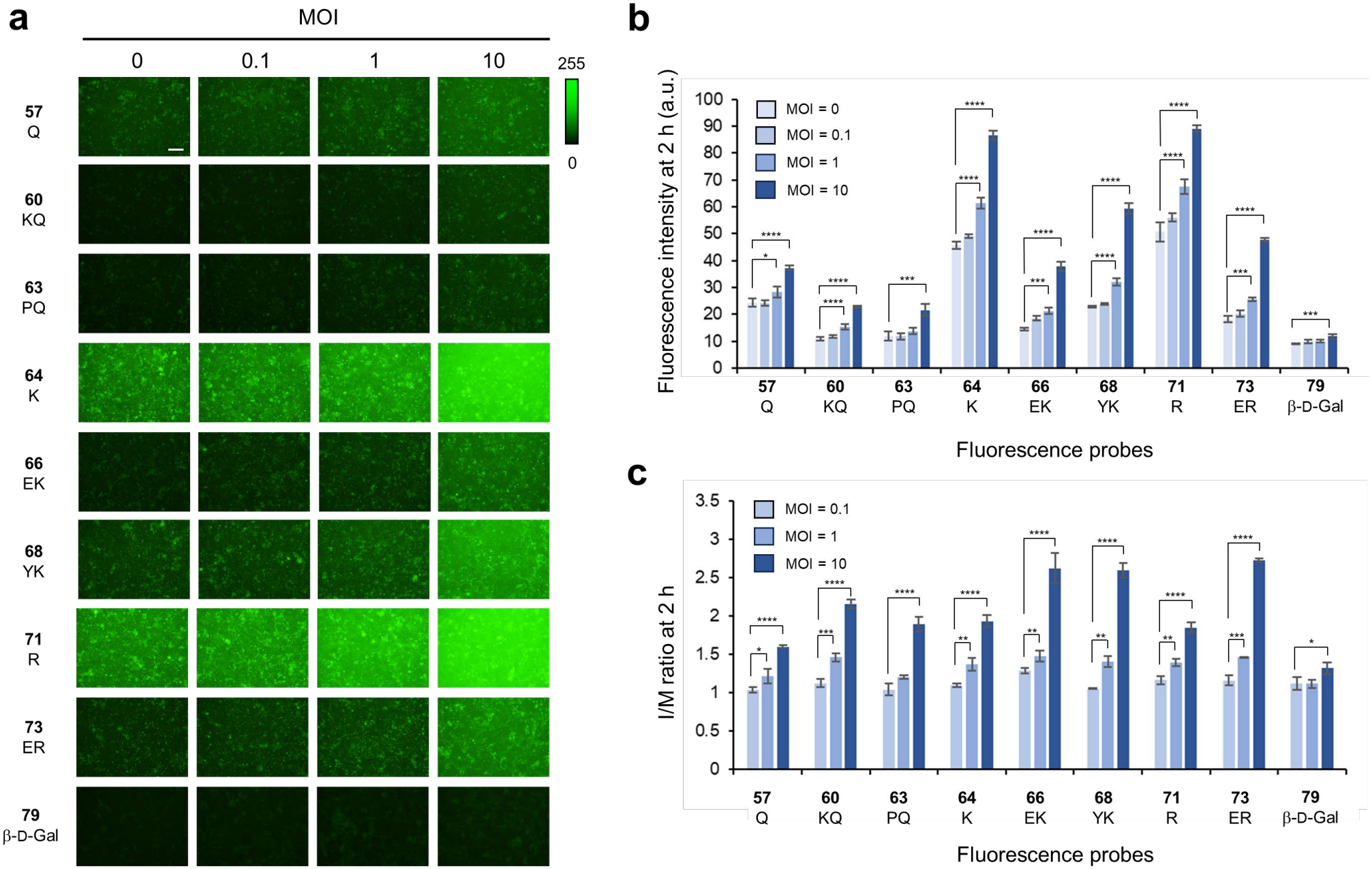
Evaluation of MOI-dependent activation of hit fluorescence probes. (**a**) Representative live-cell fluorescence images of SARS-CoV-2-infected Calu-3 cells using 9 selected hit probes at various MOIs. Calu-3 cells were infected with SARS-CoV-2 at MOI values of 0, 0.1, 1 or 10, and fluorescence probes were added at 24 h post-infection. All images were acquired 2h after the administration of each fluorescence probe. [probes] = 10 μM. Scale bar, 200 μm. (**b**) Fluorescence intensity at 2 h, quantified from live-cell fluorescence images. Error bars represent SD calculated from three images per condition (n = 3). Statistical significance was determined separately for each probe using one-way ANOVA with Dunnett’s multiple comparisons test, with MOI = 0 as the control group. Only statistically significant comparisons involving MOI = 1 or 10 are indicated (\**p*<0.05, \*\**p*<0.01, \*\*\**p*<0.001, \*\*\*\**p*<0.0001). (**c**) Ratio of fluorescence intensity in infected cells to that in mock-infected cells (I/M) at 2 h. Error bars represent SD calculated from three images per condition (n = 3). Statistical significance was determined separately for each probe using one-way ANOVA with Dunnett’s multiple comparisons test, with comparisons made against MOI = 0.1. Only statistically significant comparisons are indicated (\**p*<0.05, \*\**p*<0.01, \*\*\**p*<0.001, \*\*\*\**p*<0.0001).

### Evaluation of hit fluorescence probes in influenza virus-infected Calu-3 cells

To assess whether these probes also show enhanced activation in cells infected with other viruses, we evaluated their fluorescence responses in influenza virus-infected Calu-3 cells by live-cell imaging (**Fig. 3a**). Calu-3 cells were infected with influenza virus at an MOI of 1, and infection was confirmed by immunofluorescence staining of nucleoprotein (NP) (**Fig. S4**). In contrast to SARS-CoV-2 infection, YK-HMRG showed a slight decrease in fluorescence intensity in influenza virus-infected cells, whereas no apparent difference in fluorescence intensity was observed between mock-infected and influenza virus-infected Calu-3 cells for any of the other probes tested (**Fig. 3b, c**).

**Figure 3.**
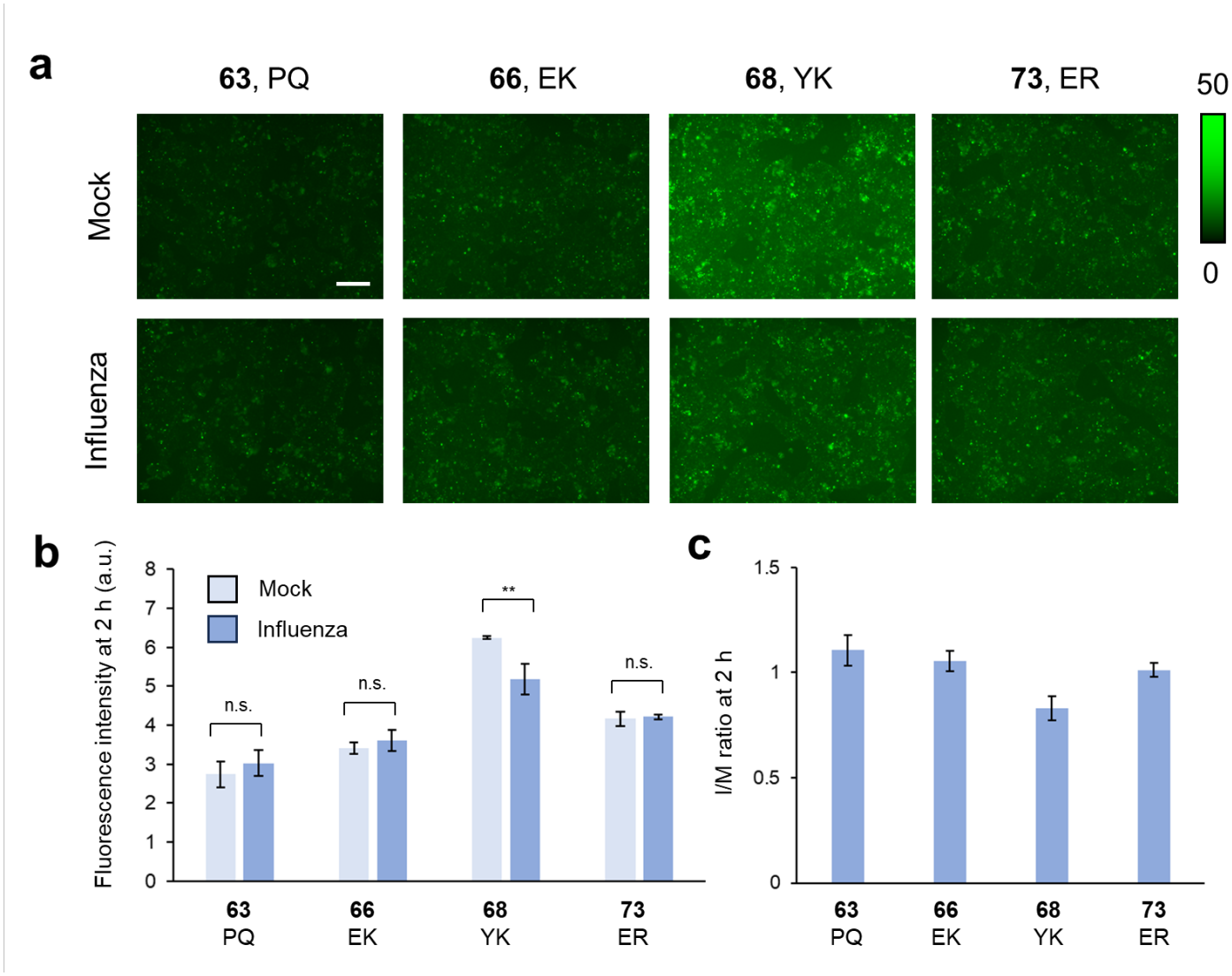
Evaluation of representative probes with influenza virus-infected Calu-3 cells. (**a**) Live-cell imaging of influenza virus-infected Calu-3 cells using representative hit fluorescence probes (PQ-HMRG, EK-HMRG, YK-HMRG and ER-HMRG). Calu-3 cells were infected with influenza virus at an MOI of 1, and fluorescence probes were added at 24 h post-infection. [probes] = 10 μM. Scale bar, 200 μm. (**b**) Fluorescence intensity of live-cell images at 2 h. Error bars represent SD calculated from three images per condition (n = 3). The significance of differences was determined by one-way ANOVA with Tukey’s HSD (n.s., not significant; \*\**p*<0.01). (**c**) I/M ratio of fluorescence intensity at 2 h. Error bars represent SD calculated from three images per condition (n = 3).

### Target identification of hit fluorescence probes

To identify the target enzymes responsible for probe activation, we selected three hit probes with different P1 amino acid residues—PQ-HMRG, YK-HMRG and ER-HMRG—based on the MOI-dependence of the fluorescence responses and their I/M ratios. We then performed diced electrophoresis gel (DEG) assay,^26^ which combines 2D gel assay with peptide mass fingerprinting analysis (**Fig. 4a**). PQ-HMRG and YK-HMRG exhibited distinct fluorescent spots, from which multiple proteins were identified. Among these, CAPN1 and cytosolic aminopeptidase (LAP3) were identified as potential target enzymes of PQ-HMRG (**Table S2**), while PSA was identified as a potential target enzyme of YK-HMRG (**Table S3**). In contrast, ER-HMRG produced multiple fluorescent spots, suggesting that multiple arginine-cleaving peptidases may contribute to its fluorescence activation.

**Figure 4.**
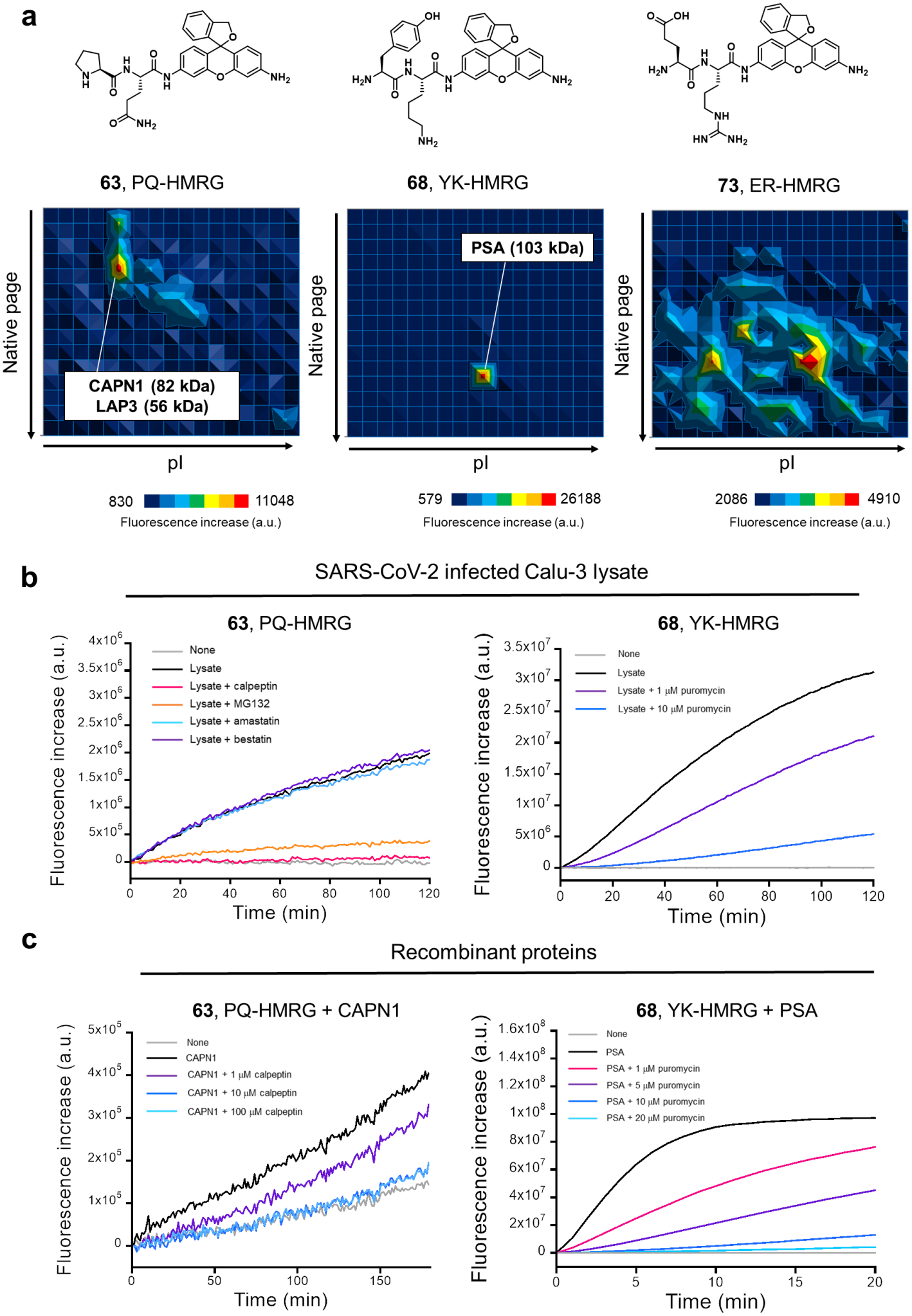
Target identification of hit probes. (**a**) Chemical structures of PQ-HMRG, YK-HMRG and ER-HMRG (upper) and identification of their target enzymes by DEG assay (bottom). CAPN1 and LAP3 were identified as target candidates for PQ-HMRG, whereas PSA was identified for YK-HMRG. Multiple fluorescence spots were observed with ER-HMRG. (**b**) Time-dependent fluorescence increases of PQ-HMRG (left) and YK-HMRG (right) in SARS-CoV-2-infected Calu-3 cell lysates in the presence or absence of enzyme inhibitors. PQ-HMRG activation was evaluated in the presence of calpeptin, MG132, amastatin or bestatin, whereas YK-HMRG activation was evaluated in the presence of puromycin. All reactions were performed in PBS. [fluorescence probe] = 1 µM, [Triton X-100] = 500 µM, [lysate] = 0.15 mg/mL, [calpeptin, MG132, amastatin, bestatin] = 10 µM, [puromycin] = 1 or 10 µM. (**c**) Time-dependent fluorescence increases of PQ-HMRG (left) and YK-HMRG (right) upon reaction with recombinant CAPN1 and PSA, respectively, in the presence or absence of their respective inhibitors. The reaction of PQ-HMRG with CAPN1 was performed in 25 mM Tris-HCl buffer (pH 7.5) containing 5 mM CaCl_2_. [PQ-HMRG] = 50 µM, [CAPN1] = 50 μg/mL, [calpeptin] = 1, 10 or 100 µM. The reaction of YK-HMRG with PSA was performed in PBS (-) containing 500 µM Triton X-100. [YK-HMRG] = 1 µM, [puromycin] = 1, 5, 10 or 20 µM, [PSA] = 10 ng/mL.

To further validate the candidate target enzymes, we first focused on PQ-HMRG and evaluated its fluorescence activation by SARS-CoV-2-infected Calu-3 cell lysates in the presence of a CAPN1 inhibitor (calpeptin or MG132) or a LAP3 inhibitor (amastatin or bestatin). The fluorescence increase of PQ-HMRG was markedly suppressed only by calpeptin and MG132, while amastatin and bestatin showed no inhibitory effect (**Fig. 4b**). These results suggested that CAPN1 is the major enzyme responsible for PQ-HMRG activation. Similarly, we evaluated the fluorescence activation of YK-HMRG in the presence of puromycin, an inhibitor of PSA. Puromycin strongly suppressed the fluorescence increase of YK-HMRG in SARS-CoV-2-infected Calu-3 cell lysates, suggesting that PSA is the major target enzyme responsible for YK-HMRG activation (**Fig. 4b**).

We further confirmed that PQ-HMRG reacts with CAPN1 recombinant protein (**Fig. 4c**). In addition, Q-HMRG showed fluorescence activation in the presence of CAPN1, suggesting that probes bearing glutamine at the P1 position are specifically activated by CAPN1 activity (**Fig. S5**). YK-HMRG was also shown to react with PSA recombinant protein, exhibiting a marked fluorescence increase (**Fig. 4c**). Furthermore, K-HMRG and EK-HMRG, which were identified as hit probes, as well as PK-HMRG, which showed a response close to the hit-selection criteria, were also activated by PSA, suggesting that probes bearing lysine at the P1 position are activated by PSA (**Fig. S6**), in accordance with our previous findings.^18, 27^ These results suggest that SARS-CoV-2 infection induces increased activities of CAPN1 and PSA in Calu-3 cells.

### PSA-dependent activation of YK-HMRG in SARS-CoV-2-infected cells

Next, we focused on YK-HMRG as a representative PSA-reactive probe that exhibits pronounced SARS-CoV-2 infection-induced fluorescence enhancement and a relatively high I/M ratio (**Fig. 2**). We performed time-lapse live-cell fluorescence imaging of mock-infected and SARS-CoV-2-infected Calu-3 cells in the presence or absence of puromycin. SARS-CoV-2-infected Calu-3 cells showed a markedly higher time-dependent fluorescence increase than mock-infected cells, suggesting that YK-HMRG can detect SARS-CoV-2-infected cells (**Fig. 5a**). Puromycin markedly suppressed YK-HMRG fluorescence activation in SARS-CoV-2-infected Calu-3 cells, indicating that the observed difference in fluorescence increase between SARS-CoV-2-infected and mock-infected Calu-3 cells is attributable to PSA activity (**Fig. 5b, c, Fig. S7**). Furthermore, we examined the relationship of PSA to SARS-CoV-2 infection by immunofluorescence staining analysis. PSA staining was observed in SARS-CoV-2-infected Calu-3 cells, particularly in regions positive for SARS-CoV-2 spike protein (**Fig. 5d, Fig. S8-9**). Cell-by-cell fluorescence intensity analysis revealed a strong positive correlation between PSA and SARS-CoV-2 spike protein signals (Pearson’s *r* = 0.824, *p* < 0.001) (**Fig. 5e, Fig. S10**). These results support the conclusion that PSA is an activity-enhanced native enzyme in SARS-CoV-2-infected Calu-3 cells.

**Figure 5.**
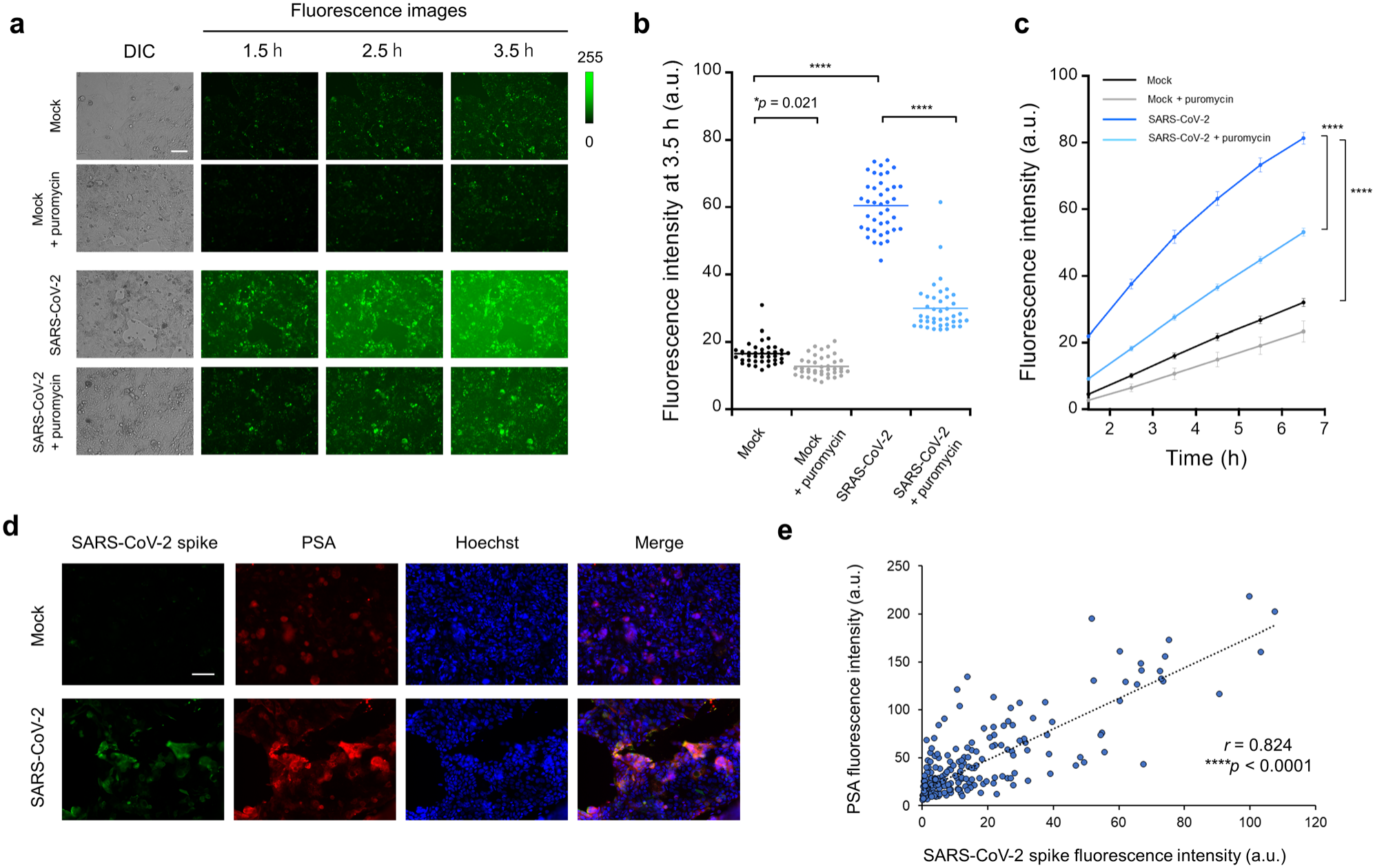
Evaluation of PSA activity and expression in SARS-CoV-2-infected Calu-3 cells. (**a**) Representative time-lapse fluorescence images of mock-infected and SARS-CoV-2-infected Calu-3 cells treated with 10 µM YK-HMRG in the presence or absence of 50 µM puromycin. Images were acquired at 1.5, 2.5, and 3.5 h after probe addition. Calu-3 cells were infected with SARS-CoV-2 at an MOI of 10, and fluorescence probes were added at 24 h post-infection. Scale bar, 200 μm. (**b**) Quantification of single-cell fluorescence intensities at 3.5 h under the four conditions shown in (**a**). Each dot represents an individual cell (n = 40 per condition), and horizontal lines indicate the mean. The significance of differences was determined by one-way ANOVA with Tukey’s HSD (\**p*<0.05, \*\*\*\**p*<0.0001). (**c**) Time-dependent changes in YK-HMRG fluorescence intensity in mock-infected and SARS-CoV-2-infected Calu-3 cells in the presence or absence of puromycin. The significance of differences was determined by one-way ANOVA with Tukey’s HSD (\*\*\*\**p*<0.0001). (**d**) Immunofluorescence staining of SARS-CoV-2 spike protein (green) and PSA (red) in mock-infected and SARS-CoV-2-infected Calu-3 cells. Calu-3 cells were infected with SARS-CoV-2 at an MOI of 10. Nuclei were stained with Hoechst (blue). Merged images are shown in the rightmost panels. Scale bar, 100 μm. (**e**) Correlation between SARS-CoV-2 spike protein and PSA fluorescence intensities. Fifty cells were randomly selected from each of five imaging fields (n = 250 cells in total), and their SARS-CoV-2 spike protein and PSA fluorescence intensities were quantified. Each dot represents a single cell, and the dotted line indicates the linear regression fit. A strong positive correlation was observed (Pearson’s *r* = 0.824, \*\*\*\**p*<0.0001).

## Discussion and conclusion

Previous transcriptomic and proteomic studies have revealed extensive alterations in host-cell gene and protein expression following SARS-CoV-2 infection.^6-8^ However, the catalytic activities of endogenous enzymes are not necessarily determined only by changes in gene or protein expression. In this study, we employed activity-based fluorescence screening using 89 enzyme-activatable fluorescent probes to investigate infection-induced changes in host-cell enzymatic activities. Screening in living Calu-3 cells identified multiple probes exhibiting enhanced fluorescence upon SARS-CoV-2 infection, including some with responses that increased in an MOI-dependent manner (**Fig. 2**). The tested hit probes were not directly activated by SARS-CoV-2 M^pro^ or PL^pro^, and several of these probes did not exhibit comparable fluorescence enhancement in influenza virus-infected Calu-3 cells under the conditions examined (**Fig. S2-3, Fig. 3**). Together, these findings suggest that the observed fluorescence responses of these probes at least partly reflect alterations in endogenous host-cell enzymatic activities associated with SARS-CoV-2 infection. However, further detailed evaluation of the enzymatic responses to other viral infections and under different infection conditions is needed to determine whether they are specific to SARS-CoV-2 infection.

Subsequent target identification, inhibitor profiling and recombinant enzyme assays identified CAPN1 and PSA as major contributors to the activation of probes containing glutamine and lysine at the P1 position, respectively, including PQ-HMRG and YK-HMRG. Among these enzymatic responses, PSA-associated activity played a particularly prominent role. Live-cell fluorescence imaging with YK-HMRG revealed marked fluorescence enhancement in SARS-CoV-2-infected cells, which was suppressed by the PSA inhibitor puromycin (**Fig. 5**). Furthermore, immunofluorescence analysis showed strong PSA staining in regions positive for SARS-CoV-2 spike protein. Together, these findings demonstrate that PSA activity is enhanced in SARS-CoV-2-infected Calu-3 cells, although the relative contributions of changes in PSA abundance and catalytic regulation remain unclear. Furthermore, compounds with CAPN1-inhibitory activity have been reported to exhibit antiviral effects,^28^ suggesting that the host enzymes identified in this study may have therapeutic relevance. This would be consistent with an analysis of SARS-CoV-2-infected individuals, which found that serum CAPN1 enzyme activity is positively correlated with the severity of pneumonia syndrome and the levels of inflammatory markers such as IL-6.^29^ Nevertheless, the specific biological roles of CAPN1 and PSA in SARS-CoV-2 infection remain to be elucidated.

Notably, a previous multi-omics study comprehensively characterized transcriptomic and proteomic changes in SARS-CoV-2-infected Calu-3 cells, the same cell model used in the present study (Pinto et al.).^7^ Interestingly, however, neither CAPN1 nor NPEPPS, the gene encoding PSA, was included among the differentially regulated transcripts or proteins reported in that study. Furthermore, examination of published translatomic and proteomic datasets from SARS-CoV-2-infected Caco-2 cells (Bojkova et al.) and transcriptomic and proteomic datasets from A549-ACE2 cells (Stukalov et al.) revealed no significant changes in NPEPPS transcript levels, translation, or protein abundance, while CAPN1 showed no significant transcriptional changes and only minor changes in protein abundance (**Table S4-5**).^6, 8^ Although there were differences in cell models and experimental conditions, these observations overall suggest that changes in enzyme activity do not necessarily correspond to changes in gene or protein expression. Our immunofluorescence staining (**Fig. 5d**) showed stronger PSA signals in SARS-CoV-2-infected cells at a high MOI (10); however, whether the enhanced enzymatic activity reflects increased PSA protein abundance or other regulatory mechanisms remains unclear. CAPN1 is a Ca^2+^-dependent protease whose activity is also regulated by post-translational modifications, while PSA is a Zn^2+^-dependent aminopeptidase whose catalytic activity is influenced by divalent metal ions.^30-34^ Such regulatory mechanisms may contribute to discrepancies between enzymatic activity and expression levels, although their involvement in the enhanced activities observed in SARS-CoV-2-infected cells remains to be investigated. In addition, enzymatic signal amplification may enable fluorescence probes to detect activity changes arising from subtle differences in enzyme abundance that may be difficult to discern through conventional expression-based analyses. Together, these findings suggest that activity-based screening can uncover functional enzymatic alterations that may not be readily identifiable through conventional multi-omics analyses, and may therefore provide a complementary perspective on host-cell responses to viral infection.

While viral enzymes have been extensively investigated as targets for infection detection and antiviral therapy,^35-37^ our findings highlight the potential value of endogenous host-cell enzymatic activities as alternative biomarkers of viral infection. The enzymatic activity information obtained through this approach may facilitate the development of novel diagnostic probes, the identification of host-directed antiviral targets, and the design of enzyme-reactive prodrugs^38, 39^ that selectively release therapeutic agents in infected cells. Although further validation in additional infection models and functional characterization of the identified enzymes are required, our findings indicate that activity-based fluorescence screening is a promising strategy for identifying candidate biomarkers and therapeutic targets in response to emerging viral threats, with potential applications extending to other infectious diseases and pathological conditions.

## Methods

### Reagents

All organic solvents and reagents were commercial products of guaranteed grade, and were used without further purification. Water was doubly distilled and deionized by a Milli-Q water system before use.

### Fluorescence probe library

HMRG and HMRef-based fluorescence probes were synthesized according to the literature.^18, 19^

### Cell lines and viruses

Calu-3 was cultured in DMEM (high glucose with L-glutamine and phenol red) and maintained at 37°C in a humidified incubator under 5% CO_2_ in air. All media were supplemented with 5% fetal bovine serum and 1% penicillin/streptomycin. Calu-3 was obtained from ATCC. The SARS-CoV-2 isolate SARS-CoV-2/Hu/DP/Kng/19-027 (provided by the Kanagawa Prefectural Institute of Public Health) and influenza A virus A/California/04/2009 (H1N1) were used in this study. Virus stocks were stored at −80 °C until use. All of the experiments on SARS-CoV-2 were performed in a biosafety level 3 containment laboratory at the Institute for Life and Medical Sciences at Kyoto University.

### Activity-based screening of fluorescence probes

Calu-3 cells (10^5^ cells) were seeded in each well of a 96-well flat plate containing 100 μL of DMEM supplemented with 5% fetal bovine serum and 1% penicillin/streptomycin and cultured overnight. Calu-3 cells were then infected with SARS-CoV-2 at an MOI of 1. At 24 h post-infection, fluorescence probes were added to the cells at a final concentration of 10 μM. Differential interference contrast (DIC) and fluorescence images were acquired immediately (0 h) and 2 h after probe addition using a CKX41 inverted fluorescence microscope (Olympus Corporation) equipped with an X-Cite 110LED illumination system (Excelitas Technologies). For each condition, three fluorescence images were acquired from different fields of view. The mean gray value of each entire image was measured using ImageJ, and FI was calculated as the average of the values obtained from the three images (n = 3 fields of view per condition). Fluorescence was detected using a B-excitation filter set (excitation, 460–490 nm; emission, ≥520 nm).

### Evaluation of MOI dependency of probe activation

Calu-3 cells (10^5^ cells) were seeded in each well of a 96-well flat plate containing 100 μL of DMEM supplemented with 5% fetal bovine serum and 1% penicillin/streptomycin and cultured overnight. Calu-3 cells were then infected with SARS-CoV-2 at an MOI of 0, 0.1, 1, or 10. At 24 h post-infection, fluorescence probes were added to the cells at a final concentration of 10 μM. DIC and fluorescence images were acquired immediately (0 h) and 2 h after probe addition as described above. The mean gray value of each entire image was measured using ImageJ, and FI was calculated as the average of the values obtained from the three images (n = 3 fields of view per condition).

### Evaluation of hit probes in influenza-infected cells

Calu-3 cells (10^5^ cells) were seeded in each well of 96-well flat plate containing 100 μL of DMEM (high glucose with L-glutamine and phenol red) supplemented with 5% fetal bovine serum and 1% penicillin/streptomycin and cultured overnight. Calu-3 cells were then infected with influenza virus at MOI of 1. At 24 h post-infection, fluorescence probes were added to the cells at a final concentration of 10 μM. Fluorescence images were acquired 2 h after probe addition using an all-in-one fluorescence microscope (BZ-X800, KEYENCE Corporation) equipped with a 10× objective lens. An 80-W metal halide lamp was used for excitation, with a GFP filter set (excitation, 470/40 nm; emission, 525/50 nm). The mean gray value of each entire image was measured using ImageJ, and FI was calculated as the average of the values obtained from the three images (n = 3 fields of view per condition). Influenza virus infection was confirmed by immunofluorescence staining (**Fig. S4**).

### DEG assay

DEG assay was performed according to the literature.^26^

### Evaluation of hit probes in SARS-CoV-2-infected Calu-3 cell lysate

SARS-CoV-2-infected Calu-3 cell lysates were prepared from cells infected with SARS-CoV-2 at an MOI of 10. At 24 h post-infection, the culture medium was removed, and CelLytic M lysis buffer (C2978, Sigma-Aldrich) was added to the cells. The cells were then collected using a cell scraper, and the resulting lysates were transferred to tubes and stored at −80 °C until use. The total protein concentrations of the cell lysates were determined using a bicinchoninic acid (BCA) protein assay. PBS (-) solutions containing 1 μM PQ-HMRG or YK-HMRG, 0.15 mg/mL SARS-CoV-2-infected Calu-3 cell lysate and 500 μM Triton X-100 with or without enzyme inhibitors were incubated for 2 h at 37 °C. For PQ-HMRG, calpeptin, MG132, amastatin or bestatin was added at a final concentration of 10 μM. For YK-HMRG, puromycin was added at a final concentration of 1 or 10 μM. Fluorescence intensity was measured over 2 h using an Envision plate reader (PerkinElmer). Filter settings were 485 (485/14 nm) for excitation and 535 (535/25 nm) for emission.

### Evaluation of reactivity of fluorescence probes with recombinant proteins

25 mM Tris-HCl buffer (pH 7.5) containing 5 mM CaCl_2_ and 50 μM PQ-HMRG and 50 μg/mL CAPN1 (208713, Sigma-Aldrich) with or without 1, 10 or 100 μM calpeptin were incubated for 3 h at 37 °C. PBS (-) solutions containing 1 μM YK-HMRG, 10 ng/mL PSA (6410-ZN, R&D Systems) and 500 μM Triton X-100 with or without 1, 5, 10 or 20 μM puromycin were incubated for 20 min at 37 °C. Fluorescence intensity was measured using an Envision plate reader (PerkinElmer). Filter settings were 485 (485/14 nm) for excitation and 535 (535/25 nm) for emission.

### Time-dependent fluorescence imaging of enzyme activity in SARS-CoV-2-infected Calu-3 cells

Calu-3 cells (10^5^ cells) were seeded in each well of a 96-well chamber containing 100 μL of DMEM supplemented with 5% fetal bovine serum and 1% penicillin/streptomycin and cultured for 24 h. Calu-3 cells were then infected with SARS-CoV-2 at an MOI of 10. At 24 h post-infection, 10 μM YK-HMRG was added with or without 50 μM puromycin. DIC and fluorescence images were acquired at 1.5, 2.5, 3.5, 4.5, 5.5 and 6.5 h after probe addition using an all-in-one fluorescence microscope (BZ-X800, KEYENCE Corporation) equipped with a 10× objective lens. For time-course analysis, the mean gray value of each entire image was measured using ImageJ, and FI was calculated as the average of the values obtained from the six images (n = 6 fields of view per condition). An 80-W metal halide lamp was used for excitation, with a GFP filter set (excitation, 470/40 nm; emission, 525/50 nm). Fluorescence intensities of individual cells in each condition were quantified using ImageJ. Regions of interest (ROIs) were manually defined for individual cells, and the mean fluorescence intensities were measured. Ten cells were randomly selected from each of four microscopic fields (n = 40 cells in total). The significance of differences was determined by one-way ANOVA with Tukey’s HSD.

### Immunofluorescence analysis

Calu-3 cells were fixed with 4% paraformaldehyde for 30 min at room temperature, permeabilized with 0.1% Triton X-100 in PBS for 10 min at room temperature, and blocked with Blocking One (Nacalai Tesque) for 30 min. After washing with PBS, the cells were incubated overnight at 4°C with mouse monoclonal anti-SARS-CoV-2 spike protein (GTX632604, GeneTex) and rabbit polyclonal anti-PSA (GTX115247, GeneTex) antibodies, each diluted 1:1,000 in PBS containing 10% Blocking One. After washing with PBS, the cells were incubated with Alexa Fluor 488-conjugated goat anti-mouse IgG (A-11001, Invitrogen) and Alexa Fluor 594-conjugated goat anti-rabbit IgG (A-11012, Invitrogen) secondary antibodies for 1 h at room temperature. Following washing with PBS, fluorescence images were acquired using an all-in-one fluorescence microscope (BZ-X800; Keyence Corporation). Fluorescence intensities of SARS-CoV-2 spike protein and PSA were quantified using ImageJ. Fifty cells were randomly selected from each of five microscopic fields (n = 250 cells in total). ROIs were manually defined for individual cells, and the mean fluorescence intensities of SARS-CoV-2 spike protein and PSA were measured within the same ROIs. Pearson’s correlation coefficients (*r*) and corresponding *p* values were calculated to evaluate the correlation between the two fluorescence signals. Linear regression lines were fitted for visualization.

## Data Availability

All data and materials are available upon request.

## Acknowledgements

This research was supported by Grants-in-Aid for JSPS Research Fellows (DC1; JP19J21205 to K.F.), Grants-in-Aid for Scientific Research (KAKENHI) (JP22K20528 to K.F.), Japan Science and Technology Agency (JST) ACT-X program (JPMJAX222G to K.F.), Masason Foundation (to K.F.), Joint Usage/Research Center Program of the Institute for Life and Medical Sciences at Kyoto University (to T.N.), the Kansai Economic Federation (KANKEIREN) (to T.N.), KAKENHI (JP19H05632 and JP24H00050 to Y.U.), JST Mirai Program (JPMJMI24G2 to Y.U.), and JST Moonshot Research and Development Program (JPMJMS2022 to Y.U.).

## Author contributions

K.F., Y.K., T.N. and Y.U. designed the project. K. F., Y.M., K.S., M.K., C.O., Y.K., T.N. and Y.U. analyzed the results. K.F. wrote the manuscript with input from all the authors. K.F., M.T. and H.W. conducted *in vitro* assay. K.F. conducted DEG assays. Y.M. and K.S. conducted live-cell imaging of infected cells and immunofluorescence analysis. The authors declare no competing interests.

## References

(1) Zhu, N.; Zhang, D.; Wang, W.; Li, X.; Yang, B.; Song, J.; Zhao, X.; Huang, B.; Shi, W.; Lu, R.; et al. A Novel Coronavirus from Patients with Pneumonia in China, 2019. New England Journal of Medicine 2020, 382 (8), 727–733. DOI: doi:10.1056/NEJMoa2001017.

(2) Hou, Y. J.; Okuda, K.; Edwards, C. E.; Martinez, D. R.; Asakura, T.; Dinnon, K. H.; Kato, T.; Lee, R. E.; Yount, B. L.; Mascenik, T. M.; et al. SARS-CoV-2 Reverse Genetics Reveals a Variable Infection Gradient in the Respiratory Tract. Cell 2020, 182 (2), 429–446.e414. DOI: 10.1016/j.cell.2020.05.042.

(3) Blanco-Melo, D.; Nilsson-Payant, B. E.; Liu, W.-C.; Uhl, S.; Hoagland, D.; Møller, R.; Jordan, T. X.; Oishi, K.; Panis, M.; Sachs, D.; et al. Imbalanced Host Response to SARS-CoV-2 Drives Development of COVID-19. Cell 2020, 181 (5), 1036–1045.e1039. DOI: 10.1016/j.cell.2020.04.026.

(4) Gordon, D. E.; Jang, G. M.; Bouhaddou, M.; Xu, J.; Obernier, K.; White, K. M.; O’Meara, M. J.; Rezelj, V. V.; Guo, J. Z.; Swaney, D. L.; et al. A SARS-CoV-2 protein interaction map reveals targets for drug repurposing. Nature 2020, 583 (7816), 459–468. DOI: 10.1038/s41586-020-2286-9.

(5) Messner, C. B.; Demichev, V.; Wendisch, D.; Michalick, L.; White, M.; Freiwald, A.; Textoris-Taube, K.; Vernardis, S. I.; Egger, A.-S.; Kreidl, M.; et al. Ultra-High-Throughput Clinical Proteomics Reveals Classifiers of COVID-19 Infection. Cell Systems 2020, 11 (1), 11–24.e14. DOI: 10.1016/j.cels.2020.05.012.

(6) Stukalov, A.; Girault, V.; Grass, V.; Karayel, O.; Bergant, V.; Urban, C.; Haas, D. A.; Huang, Y.; Oubraham, L.; Wang, A.; et al. Multilevel proteomics reveals host perturbations by SARS-CoV-2 and SARS-CoV. Nature 2021, 594 (7862), 246–252. DOI: 10.1038/s41586-021-03493-4.

(7) Pinto, S. M.; Subbannayya, Y.; Kim, H.; Hagen, L.; Górna, M. W.; Nieminen, A. I.; Bjørås, M.; Espevik, T.; Kainov, D.; Kandasamy, R. K. Multi-OMICs landscape of SARS-CoV-2-induced host responses in human lung epithelial cells. iScience 2023, 26 (1), 105895. DOI: 10.1016/j.isci.2022.105895.

(8) Bojkova, D.; Klann, K.; Koch, B.; Widera, M.; Krause, D.; Ciesek, S.; Cinatl, J.; Münch, C. Proteomics of SARS-CoV-2-infected host cells reveals therapy targets. Nature 2020, 583 (7816), 469–472. DOI: 10.1038/s41586-020-2332-7.

(9) Crozier, T.; Greenwood, E.; Williamson, J.; Guo, W.; Porter, L.; Gabaev, I.; Teixeira-Silva, A.; Grice, G.; Wickenhagen, A.; Stanton, R.; et al. Quantitative proteomic analysis of SARS-CoV-2 infection of primary human airway ciliated cells and lung epithelial cells demonstrates the effectiveness of SARS-CoV-2 innate immune evasion [version 1; peer review: 2 approved]. Wellcome Open Research 2022, 7 (224). DOI: 10.12688/wellcomeopenres.17946.1.

(10) Jin, Z.; Du, X.; Xu, Y.; Deng, Y.; Liu, M.; Zhao, Y.; Zhang, B.; Li, X.; Zhang, L.; Peng, C.; et al. Structure of Mpro from SARS-CoV-2 and discovery of its inhibitors. Nature 2020, 582 (7811), 289–293. DOI: 10.1038/s41586-020-2223-y.

(11) Shin, D.; Mukherjee, R.; Grewe, D.; Bojkova, D.; Baek, K.; Bhattacharya, A.; Schulz, L.; Widera, M.; Mehdipour, A. R.; Tascher, G.; et al. Papain-like protease regulates SARS-CoV-2 viral spread and innate immunity. Nature 2020, 587 (7835), 657–662. DOI: 10.1038/s41586-020-2601-5.

(12) Shapira, T.; Monreal, I. A.; Dion, S. P.; Buchholz, D. W.; Imbiakha, B.; Olmstead, A. D.; Jager, M.; Désilets, A.; Gao, G.; Martins, M.; et al. A TMPRSS2 inhibitor acts as a pan-SARS-CoV-2 prophylactic and therapeutic. Nature 2022, 605 (7909), 340–348. DOI: 10.1038/s41586-022-04661-w.

(13) Bojkova, D.; Reus, P.; Panosch, L.; Bechtel, M.; Rothenburger, T.; Kandler, J. D.; Pfeiffer, A.; Wagner, J. U. G.; Shumliakivska, M.; Dimmeler, S.; et al. Identification of novel antiviral drug candidates using an optimized SARS-CoV-2 phenotypic screening platform. iScience 2023, 26 (2), 105944. DOI: 10.1016/j.isci.2023.105944.

(14) Ferreira, A. C.; Soares, V. C.; de Azevedo-Quintanilha, I. G.; Dias, S. d. S. G.; Fintelman-Rodrigues, N.; Sacramento, C. Q.; Mattos, M.; de Freitas, C. S.; Temerozo, J. R.; Teixeira, L.; et al. SARS-CoV-2 engages inflammasome and pyroptosis in human primary monocytes. Cell Death Discovery 2021, 7 (1), 43. DOI: 10.1038/s41420-021-00428-w.

(15) Heal, W. P.; Dang, T. H. T.; Tate, E. W. Activity-based probes: discovering new biology and new drug targets. Chemical Society Reviews 2011, 40 (1), 246–257. DOI: 10.1039/c0cs00004c.

(16) Kozoriz, K.; Lee, J.-S. Chemical proteomics for a comprehensive understanding of functional activity and the interactome. Chemical Society Reviews 2025, 54 (13), 6186–6207. DOI: 10.1039/d5cs00381d.

(17) Fujita, K.; Urano, Y. Activity-Based Fluorescence Diagnostics for Cancer. Chemical Reviews 2024, 124 (7), 4021–4078. DOI: 10.1021/acs.chemrev.3c00612.

(18) Kuriki, Y.; Yoshioka, T.; Kamiya, M.; Komatsu, T.; Takamaru, H.; Fujita, K.; Iwaki, H.; Nanjo, A.; Akagi, Y.; Takeshita, K.; et al. Development of a fluorescent probe library enabling efficient screening of tumour-imaging probes based on discovery of biomarker enzymatic activities. Chemical Science 2022, 13 (16), 4474–4481, 10.1039/D1SC06889J. DOI: 10.1039/D1SC06889J.

(19) Fujita, K.; Kamiya, M.; Yoshioka, T.; Ogasawara, A.; Hino, R.; Kojima, R.; Ueo, H.; Urano, Y. Rapid and Accurate Visualization of Breast Tumors with a Fluorescent Probe Targeting α-Mannosidase 2C1. ACS Central Science 2020, 6 (12), 2217–2227. DOI: 10.1021/acscentsci.0c01189.

(20) Kobayashi, K.; Fujita, K.; Kojima, R.; Hino, R.; Urano, Y.; Sato, M. Rapid imaging of pulmonary metastasis from colorectal cancer with a red fluorescence probe targeting puromycin-sensitive aminopeptidase and dipeptidyl peptidase IV. Scientific Reports 2025, 15 (1), 43930. DOI: 10.1038/s41598-025-27717-z.

(21) Takahashi, R.; Ishizawa, T.; Inagaki, Y.; Tanaka, M.; Ogasawara, A.; Kuriki, Y.; Fujita, K.; Kamiya, M.; Ushiku, T.; Urano, Y.; et al. Real-Time Fluorescence Imaging to Identify Cholangiocarcinoma in the Extrahepatic Biliary Tree Using an Enzyme-Activatable Probe. Liver Cancer 2023, 1–13. DOI: 10.1159/000530645 (acccessed 7/3/2023).

(22) Sakabe, M.; Asanuma, D.; Kamiya, M.; Iwatate, R. J.; Hanaoka, K.; Terai, T.; Nagano, T.; Urano, Y. Rational Design of Highly Sensitive Fluorescence Probes for Protease and Glycosidase Based on Precisely Controlled Spirocyclization. Journal of the American Chemical Society 2013, 135 (1), 409–414. DOI: 10.1021/ja309688m.

(23) Asanuma, D.; Sakabe, M.; Kamiya, M.; Yamamoto, K.; Hiratake, J.; Ogawa, M.; Kosaka, N.; Choyke, P. L.; Nagano, T.; Kobayashi, H.; et al. Sensitive β-galactosidase-targeting fluorescence probe for visualizing small peritoneal metastatic tumours in vivo. Nature Communications 2015, 6 (1), 6463. DOI: 10.1038/ncomms7463.

(24) Hoffmann, M.; Kleine-Weber, H.; Schroeder, S.; Krüger, N.; Herrler, T.; Erichsen, S.; Schiergens, T. S.; Herrler, G.; Wu, N.-H.; Nitsche, A.; et al. SARS-CoV-2 Cell Entry Depends on ACE2 and TMPRSS2 and Is Blocked by a Clinically Proven Protease Inhibitor. Cell 2020, 181 (2), 271–280.e278. DOI: 10.1016/j.cell.2020.02.052.

(25) Rebendenne, A.; Chaves Valadão Ana, L.; Tauziet, M.; Maarifi, G.; Bonaventure, B.; McKellar, J.; Planès, R.; Nisole, S.; Arnaud-Arnould, M.; Moncorgé, O.; et al. SARS-CoV-2 Triggers an MDA-5-Dependent Interferon Response Which Is Unable To Control Replication in Lung Epithelial Cells. Journal of Virology 2021, 95 (8), 10.1128/jvi.02415-02420. DOI: 10.1128/jvi.02415-20 (acccessed 2026/11/03).

(26) Komatsu, T.; Hanaoka, K.; Adibekian, A.; Yoshioka, K.; Terai, T.; Ueno, T.; Kawaguchi, M.; Cravatt, B. F.; Nagano, T. Diced Electrophoresis Gel Assay for Screening Enzymes with Specified Activities. Journal of the American Chemical Society 2013, 135 (16), 6002–6005. DOI: 10.1021/ja401792d.

(27) Sekine, H.; Fujita, K.; Yoshida, E.; Ueno, T.; Komatsu, T.; Hayashi, T.; Ogishima, D.; Sugimura, Y.; Urano, Y.; Terao, Y. Topical Activatable Fluorescence Probes for Rapid Intraoperative Detection of Peritoneal Dissemination in High-Grade Serous Ovarian Carcinoma. bioRxiv 2026, 2026.2007.2021.739701. DOI: 10.64898/2026.07.21.739701.

(28) Xie, X.; Lan, Q.; Zhao, J.; Zhang, S.; Liu, L.; Zhang, Y.; Xu, W.; Shao, M.; Peng, J.; Xia, S.; et al. Structure-based design of pan-coronavirus inhibitors targeting host cathepsin L and calpain-1. Signal Transduction and Targeted Therapy 2024, 9 (1), 54. DOI: 10.1038/s41392-024-01758-8.

(29) Loinjak, D.; Mihić, D.; Smolić, R.; Maričić, L.; Šahinović, I.; Smolić, M.; Sikora, R.; Loinjak, S.; Dinjar, K.; Včev, A. The Correlation of Serum Calpain 1 Activity and Concentrations of Interleukin 33 in COVID-19 Acute Respiratory Distress Syndrome. Biomedicines 2023, 11 (7), 1847.

(30) Sharma, J.; Mulherkar, S.; Chen, U.-I.; Xiong, Y.; Bajaj, L.; Cho, B.-K.; Goo, Y. A.; Leung, H.-C. E.; Tolias, K. F.; Sardiello, M. Calpain activity is negatively regulated by a KCTD7–Cullin-3 complex via non-degradative ubiquitination. Cell Discovery 2023, 9 (1), 32. DOI: 10.1038/s41421-023-00533-3.

(31) Du, M.; Li, X.; Li, Z.; Shen, Q.; Wang, Y.; Li, G.; Zhang, D. Phosphorylation regulated by protein kinase A and alkaline phosphatase play positive roles in μ-calpain activity. Food Chemistry 2018, 252, 33–39. DOI: 10.1016/j.foodchem.2018.01.103.

(32) Liu, R.; Li, Y.; Wang, M.; Zhou, G.; Zhang, W. Effect of protein S-nitrosylation on autolysis and catalytic ability of μ-calpain. Food Chemistry 2016, 213, 470–477. DOI: 10.1016/j.foodchem.2016.06.104.

(33) Sengupta, S.; Horowitz, P. M.; Karsten, S. L.; Jackson, G. R.; Geschwind, D. H.; Fu, Y.; Berry, R. W.; Binder, L. I. Degradation of Tau Protein by Puromycin-Sensitive Aminopeptidase in Vitro. Biochemistry 2006, 45 (50), 15111–15119. DOI: 10.1021/bi061830d (acccessed 10/22/2026).

(34) Madabushi, S.; Chow, K. M.; Song, E. S.; Goswami, A.; Hersh, L. B.; Rodgers, D. W. Structure of puromycin-sensitive aminopeptidase and polyglutamine binding. PLOS ONE 2023, 18 (7), e0287086. DOI: 10.1371/journal.pone.0287086.

(35) Meyer, B.; Chiaravalli, J.; Gellenoncourt, S.; Brownridge, P.; Bryne, D. P.; Daly, L. A.; Grauslys, A.; Walter, M.; Agou, F.; Chakrabarti, L. A.; et al. Characterising proteolysis during SARS-CoV-2 infection identifies viral cleavage sites and cellular targets with therapeutic potential. Nature Communications 2021, 12 (1), 5553. DOI: 10.1038/s41467-021-25796-w.

(36) Rut, W.; Groborz, K.; Zhang, L.; Sun, X.; Zmudzinski, M.; Pawlik, B.; Wang, X.; Jochmans, D.; Neyts, J.; Młynarski, W.; et al. SARS-CoV-2 Mpro inhibitors and activity-based probes for patient-sample imaging. Nature chemical biology 2021, 17 (2), 222–228. DOI: 10.1038/s41589-020-00689-z.

(37) Li, G.; Hilgenfeld, R.; Whitley, R.; De Clercq, E. Therapeutic strategies for COVID-19: progress and lessons learned. Nature Reviews Drug Discovery 2023, 22 (6), 449–475. DOI: 10.1038/s41573-023-00672-y.

(38) Fujita, K.; Kamiya, M.; Dan, S.; Tachibana, R.; Kawatani, M.; Kojima, R.; Hino, R.; Kobayashi, K.; Inoue, S.; Tani, M.; et al. Enzyme activity as an actionable axis for small-molecule precision oncology. bioRxiv 2026, 2026.2004.2014.717586. DOI: 10.64898/2026.04.14.717586.

(39) Tehler, U.; Nelson, C. H.; Peterson, L. W.; Provoda, C. J.; Hilfinger, J. M.; Lee, K.-D.; McKenna, C. E.; Amidon, G. L. Puromycin-sensitive aminopeptidase: An antiviral prodrug activating enzyme. Antiviral Research 2010, 85 (3), 482–489. DOI: 10.1016/j.antiviral.2009.12.003.

