## Supplementary Information for "Activity-based fluorescence screening uncovers host enzymatic responses to SARS-CoV-2 infection"

[1]Graduate School of Medicine and [2]Graduate School of Pharmaceutical Sciences, The University of Tokyo, 7-3-1 Hongo, Bunkyo-ku, Tokyo 113-0033, Japan. [3]Division of Molecular Pharmacology, Cancer Chemotherapy Center, Japanese Foundation for Cancer Research, 3-8-31 Ariake, Koto-ku, Tokyo 135-8550, Japan. [4]Laboratory of Ultrastructural Virology, Department of Virus Research, Institute for Life and Medical Science, Kyoto University, 53 Shogoin-Kawahara-cho, Sakyo-ku, Kyoto 606-8507, Japan. [5]Laboratory of Systems Virology, Department of Biosystems Science, Institute for Life and Medical Science, Kyoto University, 53 Shogoin-Kawahara-cho, Sakyo-ku, Kyoto 606-8507, Japan. [6]Laboratory for Chemistry and Life Science, Institute of Integrated Research, Institute of Science Tokyo, Yokohama, Kanagawa 226-8501, Japan.

[†]These authors contributed equally to this work.

*

### Table of contents


**Experimental methods** · · · 3-4

**Table S1. List of fluorescence probes used in this study** · · · 5

**Figure S1. Quantitative heatmaps of fluorescence responses obtained from the activity-based screening of 89 fluorescence probes** · · · 6

**Figure S2. Evaluation of the reactivity of hit fluorescence probes with SARS-CoV-2 $M^{pro}$** · · · 7

**Figure S3. Evaluation of the reactivity of hit fluorescence probes with SARS-CoV-2 $PL^{pro}$** · · · 8

**Figure S4. Immunofluorescence detection of influenza virus nucleoprotein (NP) in Calu-3 cells** · · · 9

**Table S2. List of detected proteins in DEG assay for PQ-HMRG** · · · 10-13

**Table S3. List of detected proteins in DEG assay for YK-HMRG** · · · 14-15

**Figure S5. Evaluation of the reactivity of Q-HMRG with CAPN1** · · · 16

**Figure S6. Evaluation of the reactivity of fluorescence probes bearing lysine at the P1 position with PSA** · · · 17

**Figure S7. Representative time-lapse fluorescence images of mock-infected and SARS-CoV-2-infected Calu-3 cells treated with YK-HMRG in the presence or absence of puromycin** · · · 18

**Figure S8. Immunofluorescence analysis of PSA in SARS-CoV-2-infected Calu-3 cells** · · · 19

**Figure S9. Immunofluorescence analysis of PSA in mock-infected Calu-3 cells** · · · 20

**Figure S10. Correlation between SARS-CoV-2 spike protein and PSA fluorescence intensities in individual imaging fields** · · · 21

**Table S4. Translatomic and proteomic changes in CAPN1 and NPEPPS in SARS-CoV-2-infected Caco-2 cells** · · · 22

**Table S5. Transcriptomic and proteomic changes in CAPN1 and NPEPPS in SARS-CoV-2-infected A549-ACE2 cells** · · · 23

**References** · · · 24

**Evaluation of reactivity of fluorescence probes with SARS-CoV-2 M^pro^.** PBS (-) solutions containing 1 μM fluorescence probe (Q-HMRG, KQ-HMRG, PQ-HMRG, K-HMRG, EK-HMRG, YK-HMRG, R-HMRG, ER-HMRG or HMRef-β-D-Gal) or 20 μM Ac-Abu-Tle-Leu-Gln-AMC (3250-v, Peptide Institute Inc.), 10 μg/mL SARS-CoV-2 $M^{pro}$ (SAE0172, Sigma-Aldrich) and 500 μM Triton X-100 were incubated for 120 min at 37 °C. Fluorescence intensity was measured using an Envision plate reader (PerkinElmer). For HMRG or HMRef-based fluorescence probes, filter settings were 485 (485/14 nm) for excitation and 535 (535/25 nm) for emission. For Ac-Abu-Tle-Leu-Gln-AMC, filter settings were 355 (355/40 nm) for excitation and 460 (460/25 nm) for emission. Ac-Abu-Tle-Leu-Gln-AMC was used as a positive control.

**Evaluation of reactivity of fluorescence probes with SARS-CoV-2 $PL^{pro}$.** 50 mM HEPES-NaOH buffer (pH 7.4) containing 1 μM fluorescence probe (Q-HMRG, KQ-HMRG, PQ-HMRG, K-HMRG, EK-HMRG, YK-HMRG, R-HMRG, ER-HMRG or HMRef-β-D-Gal) or 1 μM Z-Arg-Leu-Arg-Gly-Gly-AMC (hY-P4388, MedChemExpress), 100 nM SARS-CoV-2 $PL^{pro}$ (40593-V07E, Sino Biological), 0.01% Triton X-100, 0.1 mg/mL BSA and 2 mM DTT were incubated for 120 min at 37 °C. Fluorescence intensity was measured using an Envision plate reader (PerkinElmer). For HMRG or HMRef-based fluorescence probes, filter settings were 485 (485/14 nm) for excitation and 535 (535/25 nm) for emission. For Z-Arg-Leu-Arg-Gly-Gly-AMC, filter settings were 355 (355/40 nm) for excitation and 460 (460/25 nm) for emission. Z-Arg-Leu-Arg-Gly-Gly-AMC was used as a positive control.

**Immunofluorescence staining of influenza virus nucleoprotein (NP) in infected Calu-3 cells using a mouse anti-NP antibody.** Calu-3 cells were fixed with 4% paraformaldehyde for 30 min at room temperature, permeabilized with 0.1% Triton X-100 in PBS for 10 min at room temperature, and blocked with Blocking One (Nacalai Tesque) for 30 min. After washing with PBS, the cells were incubated overnight at 4°C with mouse monoclonal anti-influenza A virus nucleoprotein antibody (ab20343, Abcam) diluted 1:1,000 in PBS containing 10% Blocking One. After washing with PBS, the cells were incubated with Alexa

Fluor 555-conjugated goat anti-mouse IgG (A-21422, Invitrogen) secondary antibody for 1 h at room temperature. Following washing with PBS, fluorescence images were acquired using an all-in-one fluorescence microscope (BZ-X800; KEYENCE Corporation).

**Evaluation of reactivity of Q-HMRG with CAPN1.** 25 mM Tris-HCl buffer (pH 7.5) containing 5 mM $CaCl_2$ and 10 μM Q-HMRG, and 0.1 mg/mL CAPN1 (208713, Sigma-Aldrich) were incubated for 180 min at 37 °C. Fluorescence intensity was measured using an Envision plate reader (PerkinElmer). Filter settings were 485 (485/14 nm) for excitation and 535 (535/25 nm) for emission.

**Evaluation of reactivity of fluorescence probes bearing lysine at the P1 position with PSA.** PBS (-) solutions containing 1 μM K-HMRG, EK-HMRG, YK-HMRG, PK-HMRG, 250 ng/mL PSA (6410-ZN, R&D Systems) and 500 μM Triton X-100 were incubated for 120 min at 37 °C. Fluorescence intensity was measured using an Envision plate reader (PerkinElmer). Filter settings were 485 (485/14 nm) for excitation and 535 (535/25 nm) for emission.

**Table S1. List of fluorescence probes used in this study.**

| Probe No | Probes | P2 | P1 |
|---|---|---|---|
| **1** | G-HMRG | - | G |
| **2** | KG-HMRG | K | G |
| **3** | LG-HMRG | L | G |
| **4** | A-HMRG | - | A |
| **5** | GA-HMRG | G | A |
| **6** | EA-HMRG | E | A |
| **7** | KA-HMRG | K | A |
| **8** | YA-HMRG | Y | A |
| **9** | LA-HMRG | L | A |
| **10** | PA-HMRG | P | A |
| **11** | V-HMRG | - | V |
| **12** | GV-HMRG | G | V |
| **13** | KV-HMRG | K | V |
| **14** | YV-HMRG | Y | V |
| **15** | LV-HMRG | L | V |
| **16** | PV-HMRG | P | V |
| **17** | L-HMRG | - | L |
| **18** | GL-HMRG | G | L |
| **19** | KL-HMRG | K | L |
| **20** | YL-HMRG | Y | L |
| **21** | LL-HMRG | L | L |
| **22** | PL-HMRG | P | L |
| **23** | I-HMRG | - | I |
| **24** | GI-HMRG | G | I |
| **25** | KI-HMRG | K | I |
| **26** | YI-HMRG | Y | I |
| **27** | LI-HMRG | L | I |
| **28** | PI-HMRG | P | I |
| **29** | M-HMRG | - | M |
| **30** | GM-HMRG | G | M |
| **31** | EM-HMRG | E | M |
| **32** | KM-HMRG | K | M |
| **33** | YM-HMRG | Y | M |
| **34** | LM-HMRG | L | M |
| **35** | PM-HMRG | P | M |
| **36** | F-HMRG | - | F |
| **37** | GF-HMRG | G | F |
| **38** | EF-HMRG | E | F |
| **39** | KF-HMRG | K | F |
| **40** | YF-HMRG | Y | F |
| **41** | LF-HMRG | L | F |
| **42** | PF-HMRG | P | F |
| **43** | Y-HMRG | - | Y |
| **44** | GY-HMRG | G | Y |
| **45** | EY-HMRG | E | Y |

| Probe No | Probes | P2 | P1 |
|---|---|---|---|
| **46** | KY-HMRG | K | Y |
| **47** | YY-HMRG | Y | Y |
| **48** | LY-HMRG | L | Y |
| **49** | PY-HMRG | P | Y |
| **50** | P-HMRG | - | P |
| **51** | GP-HMRG | G | P |
| **52** | EP-HMRG | E | P |
| **53** | KP-HMRG | K | P |
| **54** | YP-HMRG | Y | P |
| **55** | LP-HMRG | L | P |
| **56** | PP-HMRG | P | P |
| **57** | Q-HMRG | - | Q |
| **58** | GQ-HMRG | G | Q |
| **59** | EQ-HMRG | E | Q |
| **60** | KQ-HMRG | K | Q |
| **61** | YQ-HMRG | Y | Q |
| **62** | LQ-HMRG | L | Q |
| **63** | PQ-HMRG | P | Q |
| **64** | K-HMRG | - | K |
| **65** | GK-HMRG | G | K |
| **66** | EK-HMRG | E | K |
| **67** | KK-HMRG | K | K |
| **68** | YK-HMRG | Y | K |
| **69** | LK-HMRG | L | K |
| **70** | PK-HMRG | P | K |
| **71** | R-HMRG | - | R |
| **72** | GR-HMRG | G | R |
| **73** | ER-HMRG | E | R |
| **74** | KR-HMRG | K | R |
| **75** | YR-HMRG | Y | R |
| **76** | LR-HMRG | L | R |
| **77** | gGlu-HMRG | - | gGlu |
| **78** | HMRef-β-D-Glc | - | b-D-Glc |
| **79** | HMRef-β-D-Gal | - | β-D-Gal |
| **80** | HMRef-β-L-Gal | - | β-L-Gal |
| **81** | HMRef-β-D-Xyl | - | β-D-Xyl |
| **82** | HMRef-α-D-Man | - | α-D-Man |
| **83** | HMRef-β-D-Fuc | - | β-D-Fuc |
| **84** | HMRef-α-L-Fuc | - | α-L-Fuc |
| **85** | HMRef-β-L-Fuc | - | β-L-Fuc |
| **86** | HMRef-α-D-Ara | - | α-D-Ara |
| **87** | HMRef-α-L-Ara | - | α-L-Ara |
| **88** | HMRef-β-D-GlcNAc | - | β-D-GlcNAc |
| **89** | HMRef-β-D-GalNAc | - | β-D-GalNAc |

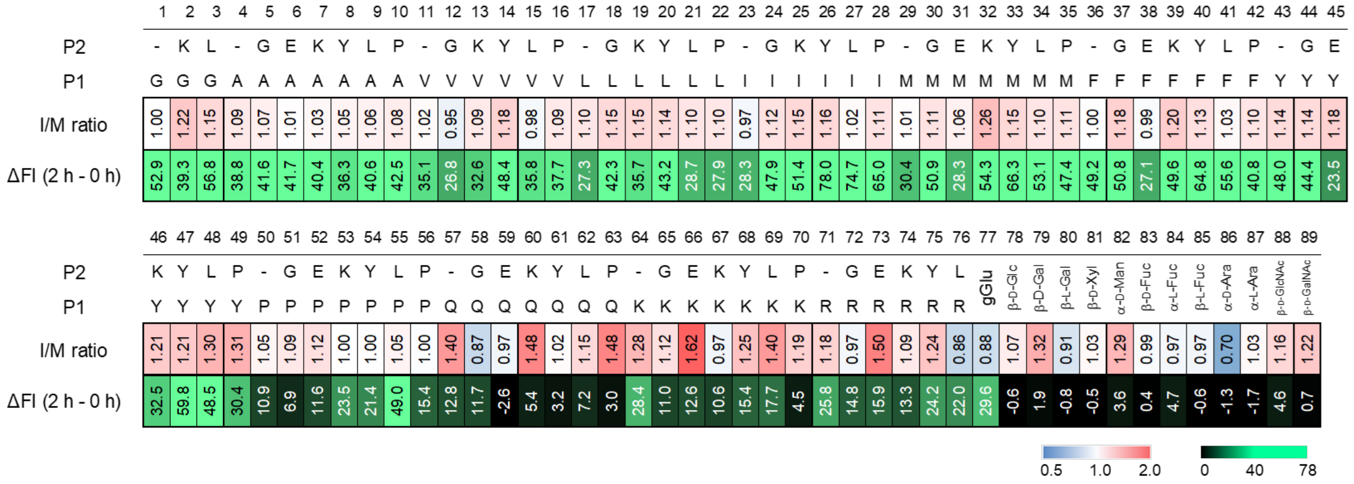


**Figure S1. Quantitative heatmaps of fluorescence responses obtained from the activity-based screening of 89 fluorescence probes.** Heatmaps showing the values of the I/M ratios at 2 h and the increases in fluorescence intensity from 0 to 2 h (ΔFI) in SARS-CoV-2-infected cells for all 89 probes shown in **Fig. 1c**. Calu-3 cells were infected with SARS-CoV-2 at a MOI of 1. Fluorescence intensities were quantified from live-cell fluorescence images acquired at 0 and 2 h after probe addition. Values represent the means of three fields of view per condition (n = 3). [Fluorescence probe] = 10 μM.

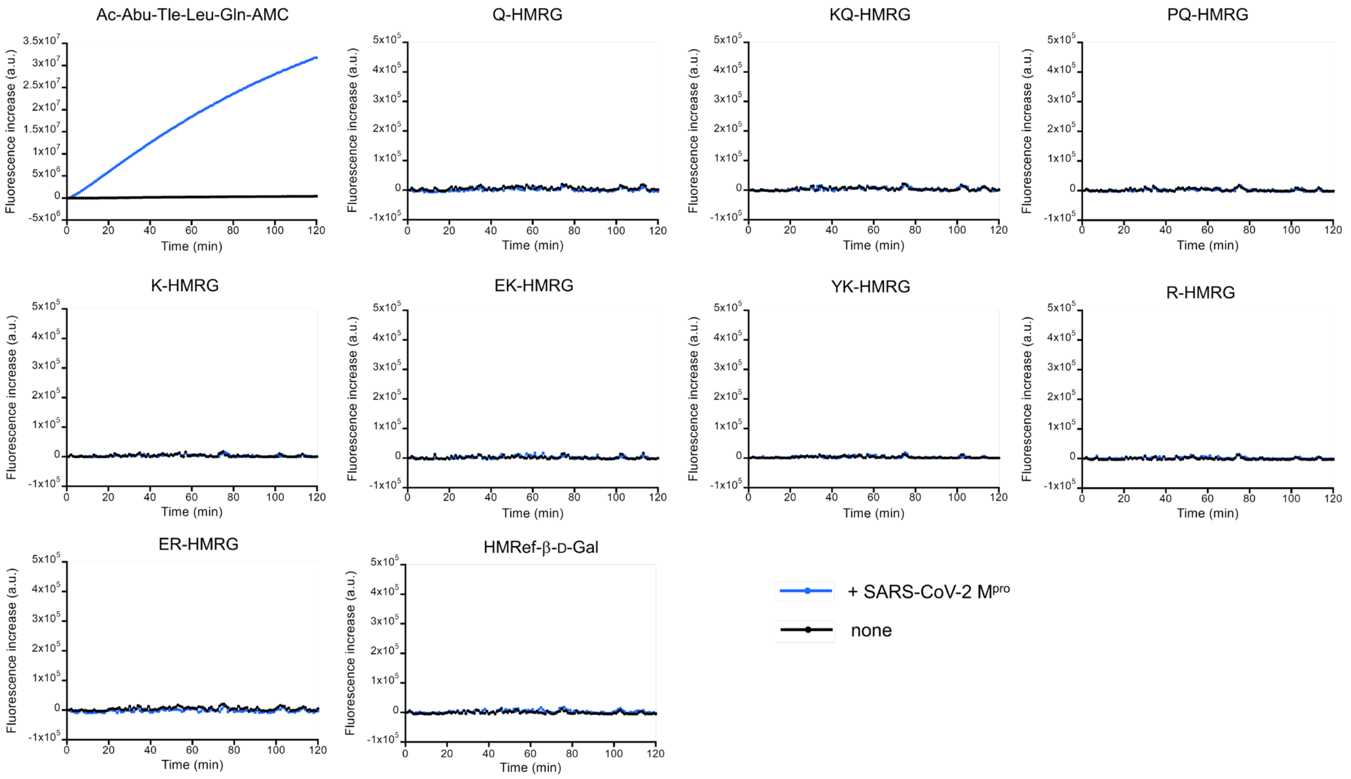


**Figure S2. Evaluation of the reactivity of hit fluorescence probes with SARS-CoV-2 Mpro.** Time-dependent fluorescence changes of the hit probes (Q-HMRG, KQ-HMRG, PQ-HMRG, K-HMRG, EK-HMRG, YK-HMRG, R-HMRG, ER-HMRG, and HMRef-β-D-Gal) in the presence or absence of SARS-CoV-2 Mpro. None of the tested fluorescence probes showed a substantial fluorescence increase in the presence of SARS-CoV-2 Mpro, whereas Ac-Abu-Tle-Leu-Gln-AMC, used as a positive control, exhibited a marked fluorescence increase. All reactions were performed in PBS (–) containing 500 μM Triton X-100 at 37 °C for 120 min. [HMRG or HMRef-based fluorescence probe] = 1 μM, [Ac-Abu-Tle-Leu-Gln-AMC] = 20 μM, and [SARS-CoV-2 Mpro] = 10 μg/mL. Fluorescence intensity was monitored using an EnVision plate reader (PerkinElmer). For HMRG- and HMRef-based fluorescence probes, filter settings were 485 (485/14 nm) for excitation and 535 (535/25 nm) for emission. For Ac-Abu-Tle-Leu-Gln-AMC, excitation and emission filter settings were 355 (355/40 nm) and 460 (460/25 nm), respectively.

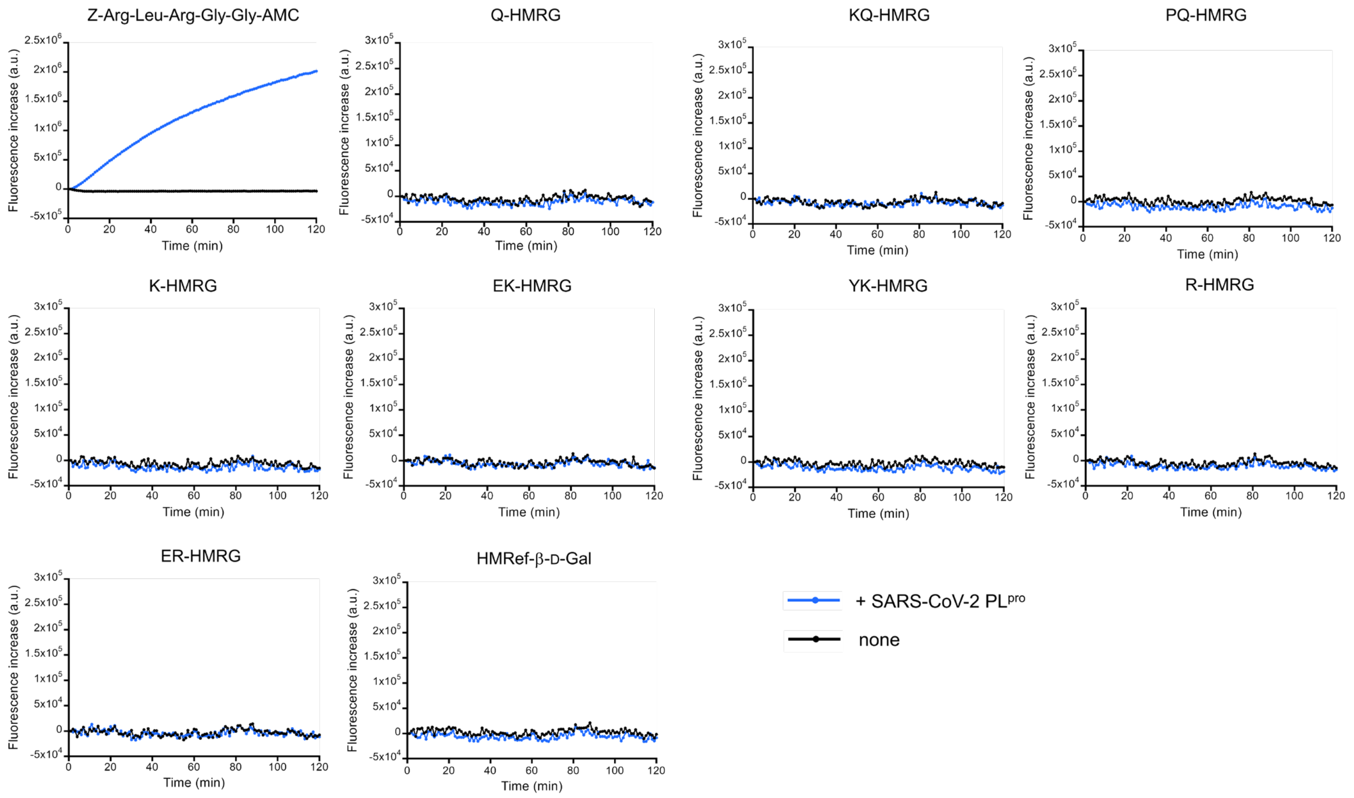


**Figure S3. Evaluation of the reactivity of hit fluorescence probes with SARS-CoV-2 PLpro.** Time-dependent fluorescence changes of the hit probes (Q-HMRG, KQ-HMRG, PQ-HMRG, K-HMRG, EK-HMRG, YK-HMRG, R-HMRG, ER-HMRG, and HMRef-β-D-Gal) in the presence or absence of SARS-CoV-2 PLpro. None of the tested fluorescence probes showed a substantial fluorescence increase in the presence of SARS-CoV-2 PLpro, whereas Z-Arg-Leu-Arg-Gly-Gly-AMC, used as a positive control, exhibited a marked fluorescence increase. All reactions were performed in 50 mM HEPES-NaOH buffer (pH7.4) containing 0.01% Triton X-100, 0.1 mg/mL BSA and 2 mM DTT at 37 °C for 120 min. [HMRG or HMRef-based fluorescence probe] = 1 μM, [Z-Arg-Leu-Arg-Gly-Gly-AMC] = 1 μM, and [SARS-CoV-2 PLpro] = 100 nM. Fluorescence intensity was monitored using an EnVision plate reader (PerkinElmer). For HMRG- and HMRef-based fluorescence probes, filter settings were 485 (485/14 nm) for excitation and 535 (535/25 nm) for emission. For Z-Arg-Leu-Arg-Gly-Gly-AMC, excitation and emission filter settings were 355 (355/40 nm) and 460 (460/25 nm), respectively.

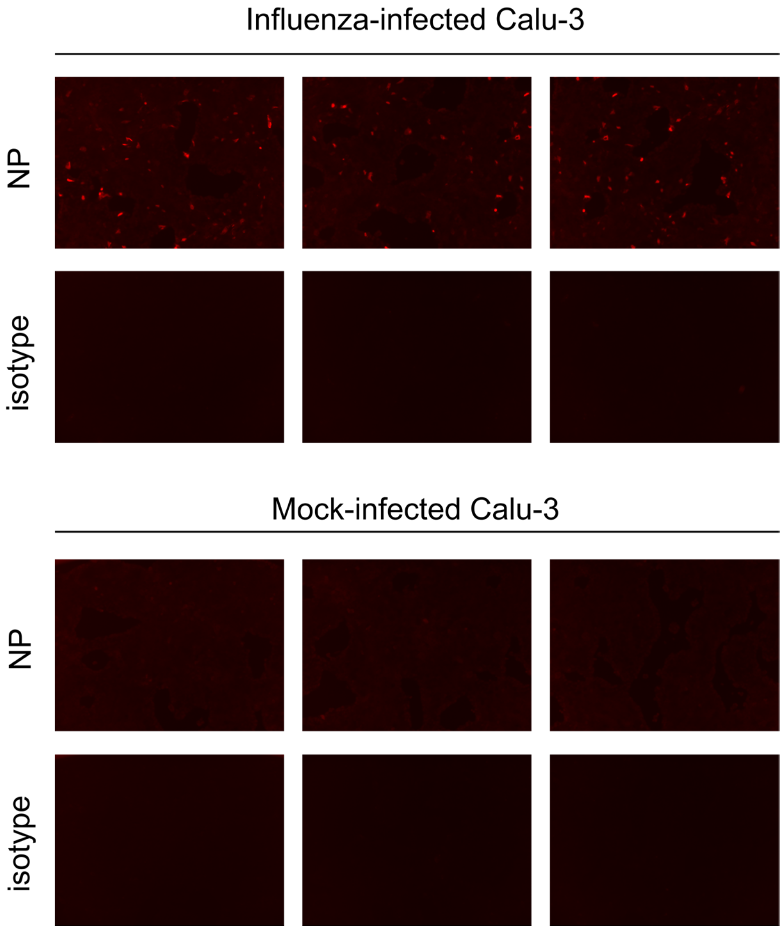


**Figure S4. Immunofluorescence detection of influenza virus nucleoprotein (NP) in Calu-3 cells.** Calu-3 cells were infected with influenza virus at a multiplicity of infection (MOI) of 1. Viral infection was confirmed by immunofluorescence staining using an anti-influenza virus NP antibody. An isotype-matched control antibody was used as a negative control to assess nonspecific antibody binding. NP-positive cells exhibited distinct red fluorescence signals, whereas minimal fluorescence was observed in the negative controls. Three fields of view were imaged for each condition (n = 3).

**Table S2. List of detected proteins in DEG assay for PQ-HMRG.** Calpain 1 (CAPN1) is highlighted in yellow. Cytosol aminopeptidase (LAP3) is highlighted in blue. 234 proteins were detected.

| | Identified Proteins (234) | ecular We | Protein Identification Probability |
|---|---|---|---|
| 1 | Keratin, type II cytoskeletal 8 OS=Homo sapiens OX=9606 GN=KRT8 PE=1 SV=7 | 54 kDa | 100% |
| 2 | Keratin, type I cytoskeletal 18 OS=Homo sapiens OX=9606 GN=KRT18 PE=1 SV=2 | 48 kDa | 100% |
| 3 | Keratin, type II cytoskeletal 1 OS=Homo sapiens OX=9606 GN=KRT1 PE=1 SV=6 | 66 kDa | 100% |
| 4 | Keratin, type I cytoskeletal 10 OS=Homo sapiens OX=9606 GN=KRT10 PE=1 SV=6 | 59 kDa | 100% |
| 5 | Keratin, type I cytoskeletal 19 OS=Homo sapiens OX=9606 GN=KRT19 PE=1 SV=4 | 44 kDa | 100% |
| 6 | Filamin-B OS=Homo sapiens OX=9606 GN=FLNB PE=1 SV=2 | 278 kDa | 100% |
| 7 | Plectin OS=Homo sapiens OX=9606 GN=PLEC PE=1 SV=3 | 532 kDa | 100% |
| 8 | Filamin-A OS=Homo sapiens OX=9606 GN=FLNA PE=1 SV=4 | 281 kDa | 100% |
| 9 | Spectrin alpha chain, non-erythrocytic 1 OS=Homo sapiens OX=9606 GN=SPTAN1 PE=1 SV=3 | 285 kDa | 100% |
| 10 | Spectrin beta chain, non-erythrocytic 1 OS=Homo sapiens OX=9606 GN=SPTBN1 PE=1 SV=2 | 275 kDa | 100% |
| 11 | Keratin, type I cytoskeletal 9 OS=Homo sapiens OX=9606 GN=KRT9 PE=1 SV=3 | 62 kDa | 100% |
| 12 | Keratin, type II cytoskeletal 2 epidermal OS=Homo sapiens OX=9606 GN=KRT2 PE=1 SV=2 | 65 kDa | 100% |
| 13 | Keratin, type II cytoskeletal 7 OS=Homo sapiens OX=9606 GN=KRT7 PE=1 SV=5 | 51 kDa | 100% |
| 14 | Talin-1 OS=Homo sapiens OX=9606 GN=TLN1 PE=1 SV=3 | 270 kDa | 100% |
| 15 | Glyceraldehyde-3-phosphate dehydrogenase OS=Homo sapiens OX=9606 GN=GAPDH PE=1 SV=3 | 36 kDa | 100% |
| 16 | Myosin-9 OS=Homo sapiens OX=9606 GN=MYH9 PE=1 SV=4 | 227 kDa | 100% |
| 17 | Heat shock 70 kDa protein 1A OS=Homo sapiens OX=9606 GN=HSPA1A PE=1 SV=1 | 70 kDa | 100% |
| 18 | Alpha-actinin-4 OS=Homo sapiens OX=9606 GN=ACTN4 PE=1 SV=2 | 105 kDa | 100% |
| 19 | Nucleolin OS=Homo sapiens OX=9606 GN=NCL PE=1 SV=3 | 77 kDa | 100% |
| 20 | Valine--tRNA ligase OS=Homo sapiens OX=9606 GN=VARS1 PE=1 SV=4 | 140 kDa | 100% |
| 21 | Serine hydroxymethyltransferase, mitochondrial OS=Homo sapiens OX=9606 GN=SHMT2 PE=1 SV=3 | 56 kDa | 100% |
| 22 | ATP-citrate synthase OS=Homo sapiens OX=9606 GN=ACLY PE=1 SV=3 | 121 kDa | 100% |
| 23 | Galectin-3-binding protein OS=Homo sapiens OX=9606 GN=LGALS3BP PE=1 SV=1 | 65 kDa | 100% |
| 24 | Actin, cytoplasmic 1 OS=Homo sapiens OX=9606 GN=ACTB PE=1 SV=1 | 42 kDa | 100% |
| 25 | ATP-dependent RNA helicase A OS=Homo sapiens OX=9606 GN=DHX9 PE=1 SV=4 | 141 kDa | 100% |
| 26 | Heat shock protein HSP 90-beta OS=Homo sapiens OX=9606 GN=HSP90AB1 PE=1 SV=4 | 83 kDa | 100% |
| 27 | Keratin, type II cytoskeletal 5 OS=Homo sapiens OX=9606 GN=KRT5 PE=1 SV=3 | 62 kDa | 100% |
| 28 | Heat shock protein HSP 90-alpha OS=Homo sapiens OX=9606 GN=HSP90AA1 PE=1 SV=5 | 85 kDa | 100% |
| 29 | Glycogen phosphorylase, brain form OS=Homo sapiens OX=9606 GN=PYGB PE=1 SV=5 | 97 kDa | 100% |
| 30 | Pyruvate kinase PKM OS=Homo sapiens OX=9606 GN=PKM PE=1 SV=4 | 58 kDa | 100% |
| 31 | SHC1_HUMAN-DECOY | | 76% |
| 32 | Interferon-induced GTP-binding protein Mx1 OS=Homo sapiens OX=9606 GN=MX1 PE=1 SV=4 | 76 kDa | 100% |
| 33 | Transitional endoplasmic reticulum ATPase OS=Homo sapiens OX=9606 GN=VCP PE=1 SV=4 | 89 kDa | 100% |
| 34 | Keratin, type II cytoskeletal 1b OS=Homo sapiens OX=9606 GN=KRT77 PE=1 SV=3 | 62 kDa | 100% |
| 35 | Heat shock cognate 71 kDa protein OS=Homo sapiens OX=9606 GN=HSPA8 PE=1 SV=1 | 71 kDa | 100% |
| 36 | Ras GTPase-activating-like protein IQGAP1 OS=Homo sapiens OX=9606 GN=IQGAP1 PE=1 SV=1 | 189 kDa | 100% |
| 37 | Fatty acid synthase OS=Homo sapiens OX=9606 GN=FASN PE=1 SV=3 | 273 kDa | 100% |
| 38 | Keratin, type I cytoskeletal 17 OS=Homo sapiens OX=9606 GN=KRT17 PE=1 SV=2 | 48 kDa | 100% |
| 39 | Ezrin OS=Homo sapiens OX=9606 GN=EZR PE=1 SV=4 | 69 kDa | 100% |
| 40 | Tubulin alpha-1B chain OS=Homo sapiens OX=9606 GN=TUBA1B PE=1 SV=1 | 50 kDa | 100% |
| 41 | Clustered mitochondria protein homolog OS=Homo sapiens OX=9606 GN=CLUH PE=1 SV=2 | 147 kDa | 100% |
| 42 | Alpha-actinin-1 OS=Homo sapiens OX=9606 GN=ACTN1 PE=1 SV=2 | 103 kDa | 100% |
| 43 | Elongation factor 2 OS=Homo sapiens OX=9606 GN=EEF2 PE=1 SV=4 | 95 kDa | 100% |
| 44 | Elongation factor 1-gamma OS=Homo sapiens OX=9606 GN=EEF1G PE=1 SV=3 | 50 kDa | 100% |
| 45 | Histidine--tRNA ligase, cytoplasmic OS=Homo sapiens OX=9606 GN=HARS1 PE=1 SV=2 | 57 kDa | 100% |
| 46 | Heterogeneous nuclear ribonucleoprotein K OS=Homo sapiens OX=9606 GN=HNRNPK PE=1 SV=1 | 51 kDa | 100% |
| 47 | Clathrin heavy chain 1 OS=Homo sapiens OX=9606 GN=CLTC PE=1 SV=5 | 192 kDa | 100% |
| 48 | Delta(3,5)-Delta(2,4)-dienoyl-CoA isomerase, mitochondrial OS=Homo sapiens OX=9606 GN=ECH1 PE=1 SV=2 | 36 kDa | 100% |
| 49 | Asparagine--tRNA ligase, cytoplasmic OS=Homo sapiens OX=9606 GN=NARS1 PE=1 SV=1 | 63 kDa | 100% |
| 50 | Fructose-bisphosphate aldolase A OS=Homo sapiens OX=9606 GN=ALDOA PE=1 SV=2 | 39 kDa | 100% |
| 51 | Receptor tyrosine-protein kinase erbB-2 OS=Homo sapiens OX=9606 GN=ERBB2 PE=1 SV=1 | 138 kDa | 100% |
| 52 | Tubulin beta chain OS=Homo sapiens OX=9606 GN=TUBB PE=1 SV=2 | 50 kDa | 100% |
| 53 | Polyadenylate-binding protein 1 OS=Homo sapiens OX=9606 GN=PABPC1 PE=1 SV=2 | 71 kDa | 100% |
| 54 | X-ray repair cross-complementing protein 5 OS=Homo sapiens OX=9606 GN=XRCC5 PE=1 SV=3 | 83 kDa | 100% |
| 55 | Eukaryotic translation initiation factor 3 subunit B OS=Homo sapiens OX=9606 GN=EIF3B PE=1 SV=3 | 92 kDa | 100% |
| 56 | Adenylyl cyclase-associated protein 1 OS=Homo sapiens OX=9606 GN=CAP1 PE=1 SV=5 | 52 kDa | 100% |
| 57 | Interleukin enhancer-binding factor 3 OS=Homo sapiens OX=9606 GN=ILF3 PE=1 SV=3 | 95 kDa | 100% |
| 58 | Cullin-3 OS=Homo sapiens OX=9606 GN=CUL3 PE=1 SV=2 | 89 kDa | 100% |
| 59 | L-lactate dehydrogenase A chain OS=Homo sapiens OX=9606 GN=LDHA PE=1 SV=2 | 37 kDa | 100% |
| 60 | Keratin, type I cytoskeletal 14 OS=Homo sapiens OX=9606 GN=KRT14 PE=1 SV=4 | 52 kDa | 100% |

| | | | |
|---|---|---|---|
| 61 | Bifunctional purine biosynthesis protein ATIC OS=Homo sapiens OX=9606 GN=ATIC PE=1 SV=3 | 65 kDa | 100% |
| 62 | Stress-induced-phosphoprotein 1 OS=Homo sapiens OX=9606 GN=STIP1 PE=1 SV=1 | 63 kDa | 100% |
| 63 | Aldehyde dehydrogenase family 1 member A3 OS=Homo sapiens OX=9606 GN=ALDH1A3 PE=1 SV=2 | 56 kDa | 100% |
| 64 | Lysine--tRNA ligase OS=Homo sapiens OX=9606 GN=KARS1 PE=1 SV=3 | 68 kDa | 100% |
| 65 | Staphylococcal nuclease domain-containing protein 1 OS=Homo sapiens OX=9606 GN=SND1 PE=1 SV=1 | 102 kDa | 100% |
| 66 | RuvB-like 1 OS=Homo sapiens OX=9606 GN=RUVBL1 PE=1 SV=1 | 50 kDa | 100% |
| 67 | Prelamin-A/C OS=Homo sapiens OX=9606 GN=LMNA PE=1 SV=1 | 74 kDa | 100% |
| 68 | 60 kDa heat shock protein, mitochondrial OS=Homo sapiens OX=9606 GN=HSPD1 PE=1 SV=2 | 61 kDa | 100% |
| 69 | Protein disulfide-isomerase A6 OS=Homo sapiens OX=9606 GN=PDIA6 PE=1 SV=1 | 48 kDa | 100% |
| 70 | General vesicular transport factor p115 OS=Homo sapiens OX=9606 GN=USO1 PE=1 SV=2 | 108 kDa | 100% |
| 71 | X-ray repair cross-complementing protein 6 OS=Homo sapiens OX=9606 GN=XRCC6 PE=1 SV=2 | 70 kDa | 100% |
| 72 | Keratin, type I cytoskeletal 13 OS=Homo sapiens OX=9606 GN=KRT13 PE=1 SV=4 | 50 kDa | 100% |
| 73 | Guanylate-binding protein 1 OS=Homo sapiens OX=9606 GN=GBP1 PE=1 SV=2 | 68 kDa | 100% |
| 74 | Eukaryotic translation initiation factor 2 subunit 3 OS=Homo sapiens OX=9606 GN=EIF2S3 PE=1 SV=3 | 51 kDa | 100% |
| 75 | Interleukin enhancer-binding factor 2 OS=Homo sapiens OX=9606 GN=ILF2 PE=1 SV=2 | 43 kDa | 100% |
| 76 | Dihydropyrimidinase-related protein 2 OS=Homo sapiens OX=9606 GN=DPYSL2 PE=1 SV=1 | 62 kDa | 100% |
| 77 | Cullin-associated NEDD8-dissociated protein 1 OS=Homo sapiens OX=9606 GN=CAND1 PE=1 SV=2 | 136 kDa | 100% |
| 78 | Isoleucine--tRNA ligase, mitochondrial OS=Homo sapiens OX=9606 GN=IARS2 PE=1 SV=2 | 114 kDa | 100% |
| 79 | Pre-mRNA-processing factor 19 OS=Homo sapiens OX=9606 GN=PRPF19 PE=1 SV=1 | 55 kDa | 100% |
| 80 | Proliferation-associated protein 2G4 OS=Homo sapiens OX=9606 GN=PA2G4 PE=1 SV=3 | 44 kDa | 100% |
| 81 | Hypoxia up-regulated protein 1 OS=Homo sapiens OX=9606 GN=HYOU1 PE=1 SV=1 | 111 kDa | 100% |
| 82 | Pre-mRNA-splicing factor ATP-dependent RNA helicase DHX15 OS=Homo sapiens OX=9606 GN=DHX15 PE=1 SV= | 91 kDa | 100% |
| 83 | Protein diaphanous homolog 1 OS=Homo sapiens OX=9606 GN=DIAPH1 PE=1 SV=2 | 141 kDa | 100% |
| 84 | Protein transport protein Sec31A OS=Homo sapiens OX=9606 GN=SEC31A PE=1 SV=3 | 133 kDa | 100% |
| 85 | Importin-7 OS=Homo sapiens OX=9606 GN=IPO7 PE=1 SV=1 | 120 kDa | 100% |
| 86 | Complement C3 OS=Homo sapiens OX=9606 GN=C3 PE=1 SV=2 | 187 kDa | 100% |
| 87 | Fibronectin OS=Homo sapiens OX=9606 GN=FN1 PE=1 SV=5 | 272 kDa | 100% |
| 88 | Albumin OS=Homo sapiens OX=9606 GN=ALB PE=1 SV=2 | 69 kDa | 100% |
| 89 | Elongation factor 1-alpha 2 OS=Homo sapiens OX=9606 GN=EEF1A2 PE=1 SV=1 | 50 kDa | 100% |
| 90 | Filaggrin-2 OS=Homo sapiens OX=9606 GN=FLG2 PE=1 SV=1 | 248 kDa | 100% |
| 91 | Vacuolar protein sorting-associated protein 35 OS=Homo sapiens OX=9606 GN=VPS35 PE=1 SV=2 | 92 kDa | 100% |
| 92 | N-alpha-acetyltransferase 15, NatA auxiliary subunit OS=Homo sapiens OX=9606 GN=NAA15 PE=1 SV=1 | 101 kDa | 100% |
| 93 | Coronin-1C OS=Homo sapiens OX=9606 GN=CORO1C PE=1 SV=1 | 53 kDa | 100% |
| 94 | Hemoglobin subunit alpha OS=Homo sapiens OX=9606 GN=HBA1 PE=1 SV=2 | 15 kDa | 100% |
| 95 | D-3-phosphoglycerate dehydrogenase OS=Homo sapiens OX=9606 GN=PHGDH PE=1 SV=4 | 57 kDa | 100% |
| 96 | AP-1 complex subunit gamma-1 OS=Homo sapiens OX=9606 GN=AP1G1 PE=1 SV=5 | 91 kDa | 100% |
| 97 | Keratin, type II cytoskeletal 6A OS=Homo sapiens OX=9606 GN=KRT6A PE=1 SV=3 | 60 kDa | 100% |
| 98 | Annexin A4 OS=Homo sapiens OX=9606 GN=ANXA4 PE=1 SV=4 | 36 kDa | 100% |
| 99 | Lysosome-associated membrane glycoprotein 2 OS=Homo sapiens OX=9606 GN=LAMP2 PE=1 SV=2 | 45 kDa | 100% |
| 100 | Endoplasmin OS=Homo sapiens OX=9606 GN=HSP90B1 PE=1 SV=1 | 92 kDa | 100% |
| 101 | Protein disulfide-isomerase A3 OS=Homo sapiens OX=9606 GN=PDIA3 PE=1 SV=4 | 57 kDa | 100% |
| 102 | Trifunctional enzyme subunit alpha, mitochondrial OS=Homo sapiens OX=9606 GN=HADHA PE=1 SV=2 | 83 kDa | 100% |
| 103 | Spliceosome RNA helicase DDX39B OS=Homo sapiens OX=9606 GN=DDX39B PE=1 SV=1 | 49 kDa | 100% |
| 104 | Importin subunit beta-1 OS=Homo sapiens OX=9606 GN=KPNB1 PE=1 SV=2 | 97 kDa | 100% |
| 105 | Heat shock protein 105 kDa OS=Homo sapiens OX=9606 GN=HSPH1 PE=1 SV=1 | 97 kDa | 100% |
| 106 | Alpha-2-macroglobulin OS=Homo sapiens OX=9606 GN=A2M PE=1 SV=3 | 163 kDa | 100% |
| 107 | Aldo-keto reductase family 1 member C3 OS=Homo sapiens OX=9606 GN=AKR1C3 PE=1 SV=4 | 37 kDa | 100% |
| 108 | Alpha-2-HS-glycoprotein OS=Homo sapiens OX=9606 GN=AHSG PE=1 SV=2 | 39 kDa | 100% |
| 109 | Glucose-6-phosphate 1-dehydrogenase OS=Homo sapiens OX=9606 GN=G6PD PE=1 SV=4 | 59 kDa | 100% |
| 110 | Hornerin OS=Homo sapiens OX=9606 GN=HRNR PE=1 SV=2 | 282 kDa | 100% |
| 111 | Caspase-14 OS=Homo sapiens OX=9606 GN=CASP14 PE=1 SV=2 | 28 kDa | 100% |
| 112 | Arginase-1 OS=Homo sapiens OX=9606 GN=ARG1 PE=1 SV=2 | 35 kDa | 100% |
| 113 | Inter-alpha-trypsin inhibitor heavy chain H2 OS=Homo sapiens OX=9606 GN=ITIH2 PE=1 SV=2 | 106 kDa | 100% |
| 114 | Protein S100-A11 OS=Homo sapiens OX=9606 GN=S100A11 PE=1 SV=2 | 12 kDa | 100% |
| 115 | Heterogeneous nuclear ribonucleoprotein U-like protein 1 OS=Homo sapiens OX=9606 GN=HNRNPUL1 PE=1 SV=2 | 96 kDa | 100% |
| 116 | RuvB-like 2 OS=Homo sapiens OX=9606 GN=RUVBL2 PE=1 SV=3 | 51 kDa | 100% |
| 117 | Heterogeneous nuclear ribonucleoprotein Q OS=Homo sapiens OX=9606 GN=SYNCRIP PE=1 SV=2 | 70 kDa | 100% |
| 118 | Cysteine--tRNA ligase, cytoplasmic OS=Homo sapiens OX=9606 GN=CARS1 PE=1 SV=3 | 85 kDa | 100% |
| 119 | Keratin, type II cytoskeletal 78 OS=Homo sapiens OX=9606 GN=KRT78 PE=1 SV=2 | 57 kDa | 100% |
| 120 | Calpain-1 catalytic subunit OS=Homo sapiens OX=9606 GN=CAPN1 PE=1 SV=1 | 82 kDa | 100% |

| | | | |
|---|---|---|---|
| 121 | Ubiquitin carboxyl-terminal hydrolase 7 OS=Homo sapiens OX=9606 GN=USP7 PE=1 SV=2 | 128 kDa | 100% |
| 122 | 26S proteasome non-ATPase regulatory subunit 2 OS=Homo sapiens OX=9606 GN=PSMD2 PE=1 SV=3 | 100 kDa | 100% |
| 123 | Mitogen-activated protein kinase 1 OS=Homo sapiens OX=9606 GN=MAPK1 PE=1 SV=3 | 41 kDa | 100% |
| 124 | Polyribonucleotide nucleotidyltransferase 1, mitochondrial OS=Homo sapiens OX=9606 GN=PNPT1 PE=1 SV=2 | 86 kDa | 100% |
| 125 | Catenin alpha-1 OS=Homo sapiens OX=9606 GN=CTNNA1 PE=1 SV=1 | 100 kDa | 100% |
| 126 | Junction plakoglobin OS=Homo sapiens OX=9606 GN=JUP PE=1 SV=3 | 82 kDa | 100% |
| 127 | Cytosol aminopeptidase OS=Homo sapiens OX=9606 GN=LAP3 PE=1 SV=3 | 56 kDa | 100% |
| 128 | AP-2 complex subunit beta OS=Homo sapiens OX=9606 GN=AP2B1 PE=1 SV=1 | 105 kDa | 100% |
| 129 | Phosphoglucomutase-1 OS=Homo sapiens OX=9606 GN=PGM1 PE=1 SV=3 | 61 kDa | 100% |
| 130 | Catalase OS=Homo sapiens OX=9606 GN=CAT PE=1 SV=3 | 60 kDa | 100% |
| 131 | Bloom syndrome protein OS=Homo sapiens OX=9606 GN=BLM PE=1 SV=1 | 159 kDa | 69% |
| 132 | ATP-dependent DNA helicase Q1 OS=Homo sapiens OX=9606 GN=RECQL PE=1 SV=3 | 73 kDa | 100% |
| 133 | Keratin, type I cytoskeletal 16 OS=Homo sapiens OX=9606 GN=KRT16 PE=1 SV=4 | 51 kDa | 100% |
| 134 | Polyadenylate-binding protein 4 OS=Homo sapiens OX=9606 GN=PABPC4 PE=1 SV=1 | 71 kDa | 100% |
| 135 | Elongation factor 1-alpha 1 OS=Homo sapiens OX=9606 GN=EEF1A1 PE=1 SV=1 | 50 kDa | 100% |
| 136 | Nascent polypeptide-associated complex subunit alpha, muscle-specific form OS=Homo sapiens OX=9606 GN=NAC | 205 kDa | 100% |
| 137 | Glutamate dehydrogenase 1, mitochondrial OS=Homo sapiens OX=9606 GN=GLUD1 PE=1 SV=2 | 61 kDa | 100% |
| 138 | Protein S100-A8 OS=Homo sapiens OX=9606 GN=S100A8 PE=1 SV=1 | 11 kDa | 100% |
| 139 | Annexin A2 OS=Homo sapiens OX=9606 GN=ANXA2 PE=1 SV=2 | 39 kDa | 100% |
| 140 | Polyubiquitin-B OS=Homo sapiens OX=9606 GN=UBB PE=1 SV=1 | 26 kDa | 100% |
| 141 | Desmoplakin OS=Homo sapiens OX=9606 GN=DSP PE=1 SV=3 | 332 kDa | 100% |
| 142 | Serine--tRNA ligase, cytoplasmic OS=Homo sapiens OX=9606 GN=SARS1 PE=1 SV=3 | 59 kDa | 100% |
| 143 | Trifunctional enzyme subunit beta, mitochondrial OS=Homo sapiens OX=9606 GN=HADHB PE=1 SV=3 | 51 kDa | 100% |
| 144 | Eukaryotic initiation factor 4A-I OS=Homo sapiens OX=9606 GN=EIF4A1 PE=1 SV=1 | 46 kDa | 100% |
| 145 | Dermcidin OS=Homo sapiens OX=9606 GN=DCD PE=1 SV=2 | 11 kDa | 100% |
| 146 | Receptor expression-enhancing protein 5 OS=Homo sapiens OX=9606 GN=REEP5 PE=1 SV=3 | 21 kDa | 100% |
| 147 | Elongation factor G, mitochondrial OS=Homo sapiens OX=9606 GN=GFM1 PE=1 SV=2 | 83 kDa | 100% |
| 148 | Protein Niban 2 OS=Homo sapiens OX=9606 GN=NIBAN2 PE=1 SV=3 | 84 kDa | 100% |
| 149 | Histone H2B type 1-K OS=Homo sapiens OX=9606 GN=H2BC12 PE=1 SV=3 | 14 kDa | 100% |
| 150 | Transcription intermediary factor 1-beta OS=Homo sapiens OX=9606 GN=TRIM28 PE=1 SV=5 | 89 kDa | 100% |
| 151 | Heterogeneous nuclear ribonucleoprotein D0 OS=Homo sapiens OX=9606 GN=HNRNPD PE=1 SV=1 | 38 kDa | 100% |
| 152 | Cofilin-1 OS=Homo sapiens OX=9606 GN=CFL1 PE=1 SV=3 | 19 kDa | 100% |
| 153 | Programmed cell death 6-interacting protein OS=Homo sapiens OX=9606 GN=PDCD6IP PE=1 SV=1 | 96 kDa | 100% |
| 154 | Enoyl-CoA hydratase, mitochondrial OS=Homo sapiens OX=9606 GN=ECHS1 PE=1 SV=4 | 31 kDa | 100% |
| 155 | Sodium/potassium-transporting ATPase subunit alpha-1 OS=Homo sapiens OX=9606 GN=ATP1A1 PE=1 SV=1 | 113 kDa | 100% |
| 156 | Keratin, type II cytoskeletal 80 OS=Homo sapiens OX=9606 GN=KRT80 PE=1 SV=2 | 51 kDa | 100% |
| 157 | 60S acidic ribosomal protein P0 OS=Homo sapiens OX=9606 GN=RPLP0 PE=1 SV=1 | 34 kDa | 100% |
| 158 | Band 4.1-like protein 2 OS=Homo sapiens OX=9606 GN=EPB41L2 PE=1 SV=1 | 113 kDa | 100% |
| 159 | Peroxiredoxin-6 OS=Homo sapiens OX=9606 GN=PRDX6 PE=1 SV=3 | 25 kDa | 100% |
| 160 | Heterogeneous nuclear ribonucleoprotein U OS=Homo sapiens OX=9606 GN=HNRNPU PE=1 SV=6 | 91 kDa | 100% |
| 161 | Protein PRRC1 OS=Homo sapiens OX=9606 GN=PRRC1 PE=1 SV=1 | 47 kDa | 100% |
| 162 | SUMO-activating enzyme subunit 2 OS=Homo sapiens OX=9606 GN=UBA2 PE=1 SV=2 | 71 kDa | 100% |
| 163 | Heterogeneous nuclear ribonucleoprotein U-like protein 2 OS=Homo sapiens OX=9606 GN=HNRNPUL2 PE=1 SV=1 | 85 kDa | 100% |
| 164 | Keratinocyte proline-rich protein OS=Homo sapiens OX=9606 GN=KPRP PE=1 SV=1 | 64 kDa | 100% |
| 165 | Nicotinamide phosphoribosyltransferase OS=Homo sapiens OX=9606 GN=NAMPT PE=1 SV=1 | 56 kDa | 100% |
| 166 | ATP-dependent DNA/RNA helicase DHX36 OS=Homo sapiens OX=9606 GN=DHX36 PE=1 SV=2 | 115 kDa | 100% |
| 167 | Argininosuccinate lyase OS=Homo sapiens OX=9606 GN=ASL PE=1 SV=4 | 52 kDa | 100% |
| 168 | Threonine--tRNA ligase 1, cytoplasmic OS=Homo sapiens OX=9606 GN=TARS1 PE=1 SV=3 | 83 kDa | 100% |
| 169 | Fumarate hydratase, mitochondrial OS=Homo sapiens OX=9606 GN=FH PE=1 SV=3 | 55 kDa | 100% |
| 170 | Protein TFG OS=Homo sapiens OX=9606 GN=TFG PE=1 SV=2 | 43 kDa | 100% |
| 171 | Pregnancy zone protein OS=Homo sapiens OX=9606 GN=PZP PE=1 SV=4 | 164 kDa | 100% |
| 172 | Exportin-7 OS=Homo sapiens OX=9606 GN=XPO7 PE=1 SV=3 | 124 kDa | 100% |
| 173 | Protein-glutamine gamma-glutamyltransferase 2 OS=Homo sapiens OX=9606 GN=TGM2 PE=1 SV=2 | 77 kDa | 100% |
| 174 | Vitronectin OS=Homo sapiens OX=9606 GN=VTN PE=1 SV=1 | 54 kDa | 98% |
| 175 | Keratin, type II cuticular Hb5 OS=Homo sapiens OX=9606 GN=KRT85 PE=1 SV=1 | 56 kDa | 97% |
| 176 | K1C9_HUMAN-DECOY | | 95% |
| 177 | 26S proteasome non-ATPase regulatory subunit 11 OS=Homo sapiens OX=9606 GN=PSMD11 PE=1 SV=3 | 47 kDa | 95% |
| 178 | HLA class I histocompatibility antigen, A alpha chain OS=Homo sapiens OX=9606 GN=HLA-A PE=1 SV=2 | 41 kDa | 95% |
| 179 | Sodium/potassium-transporting ATPase subunit beta-1 OS=Homo sapiens OX=9606 GN=ATP1B1 PE=1 SV=1 | 35 kDa | 95% |
| 180 | Alpha-enolase OS=Homo sapiens OX=9606 GN=ENO1 PE=1 SV=2 | 47 kDa | 95% |

| | | | |
|---|---|---|---|
| 181 | Inosine-5'-monophosphate dehydrogenase 2 OS=Homo sapiens OX=9606 GN=IMPDH2 PE=1 SV=2 | 56 kDa | 95% |
| 182 | Protein disulfide-isomerase A4 OS=Homo sapiens OX=9606 GN=PDIA4 PE=1 SV=2 | 73 kDa | 95% |
| 183 | Myosin light chain 6B OS=Homo sapiens OX=9606 GN=MYL6B PE=1 SV=1 | 23 kDa | 95% |
| 184 | Heterogeneous nuclear ribonucleoprotein L OS=Homo sapiens OX=9606 GN=HNRNPL PE=1 SV=2 | 64 kDa | 95% |
| 185 | CD44 antigen OS=Homo sapiens OX=9606 GN=CD44 PE=1 SV=3 | 82 kDa | 95% |
| 186 | DNA ligase 1 OS=Homo sapiens OX=9606 GN=LIG1 PE=1 SV=1 | 102 kDa | 95% |
| 187 | Inter-alpha-trypsin inhibitor heavy chain H1 OS=Homo sapiens OX=9606 GN=ITIH1 PE=1 SV=3 | 101 kDa | 95% |
| 188 | Transcription factor BTF3 OS=Homo sapiens OX=9606 GN=BTF3 PE=1 SV=1 | 22 kDa | 95% |
| 189 | Adenosylhomocysteinase OS=Homo sapiens OX=9606 GN=AHCY PE=1 SV=4 | 48 kDa | 95% |
| 190 | ATP synthase subunit alpha, mitochondrial OS=Homo sapiens OX=9606 GN=ATP5F1A PE=1 SV=1 | 60 kDa | 95% |
| 191 | Calnexin OS=Homo sapiens OX=9606 GN=CANX PE=1 SV=2 | 68 kDa | 95% |
| 192 | Elongation factor 1-delta OS=Homo sapiens OX=9606 GN=EEF1D PE=1 SV=5 | 31 kDa | 95% |
| 193 | Lon protease homolog, mitochondrial OS=Homo sapiens OX=9606 GN=LONP1 PE=1 SV=2 | 106 kDa | 95% |
| 194 | Transaldolase OS=Homo sapiens OX=9606 GN=TALDO1 PE=1 SV=2 | 38 kDa | 95% |
| 195 | Signal transducer and activator of transcription 1-alpha/beta OS=Homo sapiens OX=9606 GN=STAT1 PE=1 SV=2 | 87 kDa | 95% |
| 196 | Large proline-rich protein BAG6 OS=Homo sapiens OX=9606 GN=BAG6 PE=1 SV=2 | 119 kDa | 95% |
| 197 | T-complex protein 1 subunit gamma OS=Homo sapiens OX=9606 GN=CCT3 PE=1 SV=4 | 61 kDa | 95% |
| 198 | Hsc70-interacting protein OS=Homo sapiens OX=9606 GN=ST13 PE=1 SV=2 | 41 kDa | 95% |
| 199 | Importin subunit alpha-1 OS=Homo sapiens OX=9606 GN=KPNA2 PE=1 SV=1 | 58 kDa | 95% |
| 200 | Eukaryotic translation initiation factor 5 OS=Homo sapiens OX=9606 GN=EIF5 PE=1 SV=2 | 49 kDa | 95% |
| 201 | Small nuclear ribonucleoprotein Sm D2 OS=Homo sapiens OX=9606 GN=SNRPD2 PE=1 SV=1 | 14 kDa | 95% |
| 202 | Nodal modulator 3 OS=Homo sapiens OX=9606 GN=NOMO3 PE=1 SV=2 | 134 kDa | 95% |
| 203 | T-complex protein 1 subunit beta OS=Homo sapiens OX=9606 GN=CCT2 PE=1 SV=4 | 57 kDa | 95% |
| 204 | Desmoglein-1 OS=Homo sapiens OX=9606 GN=DSG1 PE=1 SV=2 | 114 kDa | 95% |
| 205 | Kinesin-like protein KIF1A OS=Homo sapiens OX=9606 GN=KIF1A PE=1 SV=2 | 191 kDa | 95% |
| 206 | Eukaryotic translation initiation factor 3 subunit A OS=Homo sapiens OX=9606 GN=EIF3A PE=1 SV=1 | 167 kDa | 95% |
| 207 | ATP-dependent RNA helicase DDX1 OS=Homo sapiens OX=9606 GN=DDX1 PE=1 SV=2 | 82 kDa | 95% |
| 208 | Serpin B12 OS=Homo sapiens OX=9606 GN=SERPINB12 PE=1 SV=1 | 46 kDa | 95% |
| 209 | Coronin-1B OS=Homo sapiens OX=9606 GN=CORO1B PE=1 SV=1 | 54 kDa | 95% |
| 210 | Echinoderm microtubule-associated protein-like 4 OS=Homo sapiens OX=9606 GN=EML4 PE=1 SV=3 | 109 kDa | 95% |
| 211 | Reticulon-4 OS=Homo sapiens OX=9606 GN=RTN4 PE=1 SV=2 | 130 kDa | 95% |
| 212 | COP9 signalosome complex subunit 3 OS=Homo sapiens OX=9606 GN=COPS3 PE=1 SV=3 | 48 kDa | 95% |
| 213 | 26S proteasome non-ATPase regulatory subunit 1 OS=Homo sapiens OX=9606 GN=PSMD1 PE=1 SV=2 | 106 kDa | 94% |
| 214 | Proteasome subunit alpha type-1 OS=Homo sapiens OX=9606 GN=PSMA1 PE=1 SV=1 | 30 kDa | 84% |
| 215 | DNA replication licensing factor MCM2 OS=Homo sapiens OX=9606 GN=MCM2 PE=1 SV=4 | 102 kDa | 84% |
| 216 | Skin-specific protein 32 OS=Homo sapiens OX=9606 GN=XP32 PE=1 SV=1 | 26 kDa | 73% |
| 217 | CDKAL_HUMAN-DECOY | | 68% |
| 218 | ADP-ribosylation factor 3 OS=Homo sapiens OX=9606 GN=ARF3 PE=1 SV=2 | 21 kDa | 62% |
| 219 | Arf-GAP with GTPase, ANK repeat and PH domain-containing protein 2 OS=Homo sapiens OX=9606 GN=AGAP2 PE | 125 kDa | 57% |
| 220 | Keratin, type I cytoskeletal 23 OS=Homo sapiens OX=9606 GN=KRT23 PE=1 SV=2 | 48 kDa | 53% |
| 221 | Endoplasmic reticulum chaperone BiP OS=Homo sapiens OX=9606 GN=HSPA5 PE=1 SV=2 | 72 kDa | 52% |
| 222 | Spectrin beta chain, non-erythrocytic 4 OS=Homo sapiens OX=9606 GN=SPTBN4 PE=1 SV=2 | 289 kDa | 43% |
| 223 | Adenosine deaminase-like protein OS=Homo sapiens OX=9606 GN=ADAL PE=1 SV=2 | 40 kDa | 32% |
| 224 | Histone deacetylase 7 OS=Homo sapiens OX=9606 GN=HDAC7 PE=1 SV=2 | 103 kDa | 32% |
| 225 | Protocadherin gamma-B3 OS=Homo sapiens OX=9606 GN=PCDHGB3 PE=1 SV=4 | 101 kDa | 32% |
| 226 | Desmin OS=Homo sapiens OX=9606 GN=DES PE=1 SV=3 | 54 kDa | 24% |
| 227 | Keratin, type II cytoskeletal 71 OS=Homo sapiens OX=9606 GN=KRT71 PE=1 SV=3 | 57 kDa | 8% |
| 228 | CFA74_HUMAN-DECOY | | 5% |
| 229 | Methionine aminopeptidase 2 OS=Homo sapiens OX=9606 GN=METAP2 PE=1 SV=1 | 53 kDa | 5% |
| 230 | Activated RNA polymerase II transcriptional coactivator p15 OS=Homo sapiens OX=9606 GN=SUB1 PE=1 SV=3 | 14 kDa | 5% |
| 231 | Exostosin-1 OS=Homo sapiens OX=9606 GN=EXT1 PE=1 SV=2 | 86 kDa | 5% |
| 232 | Protrudin OS=Homo sapiens OX=9606 GN=ZFYVE27 PE=1 SV=1 | 46 kDa | 5% |
| 233 | InaD-like protein OS=Homo sapiens OX=9606 GN=PATJ PE=1 SV=3 | 196 kDa | 5% |
| 234 | Spectrin beta chain, non-erythrocytic 2 OS=Homo sapiens OX=9606 GN=SPTBN2 PE=1 SV=3 | 271 kDa | 1% |

**Table S3. List of detected proteins in DEG assay for YK-HMRG.** Puromycin-sensitive aminopeptidase (PSA) is highlighted in yellow. 118 proteins were detected.

| | Identified Proteins (118) | Molecular Weight | Protein Identification Probability |
|---|---|---|---|
| 1 | Keratin, type II cytoskeletal 1 OS=Homo sapiens OX=9606 GN=KRT1 PE=1 SV=6 | 66 kDa | 100% |
| 2 | Keratin, type I cytoskeletal 10 OS=Homo sapiens OX=9606 GN=KRT10 PE=1 SV=6 | 59 kDa | 100% |
| 3 | Keratin, type I cytoskeletal 9 OS=Homo sapiens OX=9606 GN=KRT9 PE=1 SV=3 | 62 kDa | 100% |
| 4 | Keratin, type II cytoskeletal 2 epidermal OS=Homo sapiens OX=9606 GN=KRT2 PE=1 SV=2 | 65 kDa | 100% |
| 5 | Puromycin-sensitive aminopeptidase OS=Homo sapiens OX=9606 GN=NPEPPS PE=1 SV=2 | 103 kDa | 100% |
| 6 | Endoplasmic reticulum chaperone BiP OS=Homo sapiens OX=9606 GN=HSPA5 PE=1 SV=2 | 72 kDa | 100% |
| 7 | Keratin, type II cytoskeletal 5 OS=Homo sapiens OX=9606 GN=KRT5 PE=1 SV=3 | 62 kDa | 100% |
| 8 | Actin, cytoplasmic 1 OS=Homo sapiens OX=9606 GN=ACTB PE=1 SV=1 | 42 kDa | 100% |
| 9 | Keratin, type I cytoskeletal 14 OS=Homo sapiens OX=9606 GN=KRT14 PE=1 SV=4 | 52 kDa | 100% |
| 10 | Desmoglein-1 OS=Homo sapiens OX=9606 GN=DSG1 PE=1 SV=2 | 114 kDa | 100% |
| 11 | Neutral alpha-glucosidase AB OS=Homo sapiens OX=9606 GN=GANAB PE=1 SV=3 | 107 kDa | 100% |
| 12 | Albumin OS=Homo sapiens OX=9606 GN=ALB PE=1 SV=2 | 69 kDa | 100% |
| 13 | Ubiquitin-like modifier-activating enzyme 1 OS=Homo sapiens OX=9606 GN=UBA1 PE=1 SV=3 | 118 kDa | 100% |
| 14 | SHC1_HUMAN-DECOY | ? | 46% |
| 15 | Desmoplakin OS=Homo sapiens OX=9606 GN=DSP PE=1 SV=3 | 332 kDa | 100% |
| 16 | Vitamin D-binding protein OS=Homo sapiens OX=9606 GN=GC PE=1 SV=2 | 53 kDa | 100% |
| 17 | Glyceraldehyde-3-phosphate dehydrogenase OS=Homo sapiens OX=9606 GN=GAPDH PE=1 SV=3 | 36 kDa | 100% |
| 18 | Junction plakoglobin OS=Homo sapiens OX=9606 GN=JUP PE=1 SV=3 | 82 kDa | 100% |
| 19 | Guanine deaminase OS=Homo sapiens OX=9606 GN=GDA PE=1 SV=1 | 51 kDa | 100% |
| 20 | Hornerin OS=Homo sapiens OX=9606 GN=HRNR PE=1 SV=2 | 282 kDa | 100% |
| 21 | Alpha-fetoprotein OS=Homo sapiens OX=9606 GN=AFP PE=1 SV=1 | 69 kDa | 100% |
| 22 | Heat shock protein HSP 90-beta OS=Homo sapiens OX=9606 GN=HSP90AB1 PE=1 SV=4 | 83 kDa | 100% |
| 23 | Chloride intracellular channel protein 1 OS=Homo sapiens OX=9606 GN=CLIC1 PE=1 SV=4 | 27 kDa | 100% |
| 24 | Protein disulfide-isomerase OS=Homo sapiens OX=9606 GN=P4HB PE=1 SV=3 | 57 kDa | 100% |
| 25 | Leukotriene A-4 hydrolase OS=Homo sapiens OX=9606 GN=LTA4H PE=1 SV=2 | 69 kDa | 100% |
| 26 | Keratinocyte proline-rich protein OS=Homo sapiens OX=9606 GN=KPRP PE=1 SV=1 | 64 kDa | 100% |
| 27 | Antithrombin-III OS=Homo sapiens OX=9606 GN=SERPINC1 PE=1 SV=1 | 53 kDa | 100% |
| 28 | Arginase-1 OS=Homo sapiens OX=9606 GN=ARG1 PE=1 SV=2 | 35 kDa | 100% |
| 29 | Heat shock protein HSP 90-alpha OS=Homo sapiens OX=9606 GN=HSP90AA1 PE=1 SV=5 | 85 kDa | 100% |
| 30 | Glutathione S-transferase P OS=Homo sapiens OX=9606 GN=GSTP1 PE=1 SV=2 | 23 kDa | 100% |
| 31 | Translin OS=Homo sapiens OX=9606 GN=TSN PE=1 SV=1 | 26 kDa | 100% |
| 32 | Caspase-14 OS=Homo sapiens OX=9606 GN=CASP14 PE=1 SV=2 | 28 kDa | 100% |
| 33 | Heat shock 70 kDa protein 4 OS=Homo sapiens OX=9606 GN=HSPA4 PE=1 SV=4 | 94 kDa | 100% |
| 34 | Importin subunit beta-1 OS=Homo sapiens OX=9606 GN=KPNB1 PE=1 SV=2 | 97 kDa | 100% |
| 35 | Filaggrin-2 OS=Homo sapiens OX=9606 GN=FLG2 PE=1 SV=1 | 248 kDa | 100% |
| 36 | Keratin, type II cytoskeletal 78 OS=Homo sapiens OX=9606 GN=KRT78 PE=1 SV=2 | 57 kDa | 100% |
| 37 | Immunity-related GTPase family Q protein OS=Homo sapiens OX=9606 GN=IRGQ PE=1 SV=1 | 63 kDa | 100% |
| 38 | Alpha-2-HS-glycoprotein OS=Homo sapiens OX=9606 GN=AHSG PE=1 SV=2 | 39 kDa | 100% |
| 39 | Annexin A2 OS=Homo sapiens OX=9606 GN=ANXA2 PE=1 SV=2 | 39 kDa | 100% |
| 40 | Inter-alpha-trypsin inhibitor heavy chain H2 OS=Homo sapiens OX=9606 GN=ITIH2 PE=1 SV=2 | 106 kDa | 100% |
| 41 | Hsc70-interacting protein OS=Homo sapiens OX=9606 GN=ST13 PE=1 SV=2 | 41 kDa | 100% |
| 42 | Desmocollin-1 OS=Homo sapiens OX=9606 GN=DSC1 PE=1 SV=2 | 100 kDa | 100% |
| 43 | Bleomycin hydrolase OS=Homo sapiens OX=9606 GN=BLMH PE=1 SV=1 | 53 kDa | 100% |
| 44 | Calpain-2 catalytic subunit OS=Homo sapiens OX=9606 GN=CAPN2 PE=1 SV=6 | 80 kDa | 100% |
| 45 | Filaggrin OS=Homo sapiens OX=9606 GN=FLG PE=1 SV=3 | 435 kDa | 100% |
| 46 | Thimet oligopeptidase OS=Homo sapiens OX=9606 GN=THOP1 PE=1 SV=2 | 79 kDa | 100% |
| 47 | Citrate synthase, mitochondrial OS=Homo sapiens OX=9606 GN=CS PE=1 SV=2 | 52 kDa | 100% |
| 48 | Peroxiredoxin-2 OS=Homo sapiens OX=9606 GN=PRDX2 PE=1 SV=5 | 22 kDa | 100% |
| 49 | Heat shock cognate 71 kDa protein OS=Homo sapiens OX=9606 GN=HSPA8 PE=1 SV=1 | 71 kDa | 100% |
| 50 | Collagen alpha-1(III) chain OS=Homo sapiens OX=9606 GN=COL3A1 PE=1 SV=4 | 139 kDa | 100% |
| 51 | Inter-alpha-trypsin inhibitor heavy chain H3 OS=Homo sapiens OX=9606 GN=ITIH3 PE=1 SV=2 | 100 kDa | 100% |
| 52 | ATP synthase subunit beta, mitochondrial OS=Homo sapiens OX=9606 GN=ATP5F1B PE=1 SV=3 | 57 kDa | 100% |
| 53 | Plakophilin-1 OS=Homo sapiens OX=9606 GN=PKP1 PE=1 SV=2 | 83 kDa | 100% |
| 54 | Inter-alpha-trypsin inhibitor heavy chain H4 OS=Homo sapiens OX=9606 GN=ITIH4 PE=1 SV=4 | 103 kDa | 100% |
| 55 | Gamma-glutamylcyclotransferase OS=Homo sapiens OX=9606 GN=GGCT PE=1 SV=1 | 21 kDa | 100% |
| 56 | ASB6_HUMAN-DECOY | ? | 96% |
| 57 | Lactotransferrin OS=Homo sapiens OX=9606 GN=LTF PE=1 SV=6 | 78 kDa | 100% |
| 58 | Lipocalin-1 OS=Homo sapiens OX=9606 GN=LCN1 PE=1 SV=1 | 19 kDa | 100% |

| | | | |
|---|---|---|---|
| 59 | Dermcidin OS=Homo sapiens OX=9606 GN=DCD PE=1 SV=2 | 11 kDa | 100% |
| 60 | Calmodulin-like protein 5 OS=Homo sapiens OX=9606 GN=CALML5 PE=1 SV=2 | 16 kDa | 100% |
| 61 | Cystatin-A OS=Homo sapiens OX=9606 GN=CSTA PE=1 SV=1 | 11 kDa | 100% |
| 62 | Protein-glutamine gamma-glutamyltransferase E OS=Homo sapiens OX=9606 GN=TGM3 PE=1 SV=4 | 77 kDa | 100% |
| 63 | Actin, alpha cardiac muscle 1 OS=Homo sapiens OX=9606 GN=ACTC1 PE=1 SV=1 | 42 kDa | 100% |
| 64 | L-lactate dehydrogenase A chain OS=Homo sapiens OX=9606 GN=LDHA PE=1 SV=2 | 37 kDa | 100% |
| 65 | Histidine ammonia-lyase OS=Homo sapiens OX=9606 GN=HAL PE=1 SV=1 | 73 kDa | 100% |
| 66 | ATP synthase subunit alpha, mitochondrial OS=Homo sapiens OX=9606 GN=ATP5F1A PE=1 SV=1 | 60 kDa | 99% |
| 67 | Keratin, type II cytoskeletal 1b OS=Homo sapiens OX=9606 GN=KRT77 PE=1 SV=3 | 62 kDa | 99% |
| 68 | Catalase OS=Homo sapiens OX=9606 GN=CAT PE=1 SV=3 | 60 kDa | 99% |
| 69 | Protein-glutamine gamma-glutamyltransferase 2 OS=Homo sapiens OX=9606 GN=TGM2 PE=1 SV=2 | 77 kDa | 98% |
| 70 | K1C9_HUMAN-DECOY | ? | 96% |
| 71 | Afamin OS=Homo sapiens OX=9606 GN=AFM PE=1 SV=1 | 69 kDa | 95% |
| 72 | Keratin, type II cytoskeletal 80 OS=Homo sapiens OX=9606 GN=KRT80 PE=1 SV=2 | 51 kDa | 77% |
| 73 | Keratin, type II cytoskeletal 6A OS=Homo sapiens OX=9606 GN=KRT6A PE=1 SV=3 | 60 kDa | 100% |
| 74 | Keratin, type I cytoskeletal 15 OS=Homo sapiens OX=9606 GN=KRT15 PE=1 SV=3 | 49 kDa | 98% |
| 75 | Keratin, type II cytoskeletal 8 OS=Homo sapiens OX=9606 GN=KRT8 PE=1 SV=7 | 54 kDa | 97% |
| 76 | Isocitrate dehydrogenase [NADP] cytoplasmic OS=Homo sapiens OX=9606 GN=IDH1 PE=1 SV=2 | 47 kDa | 96% |
| 77 | 60S acidic ribosomal protein P0 OS=Homo sapiens OX=9606 GN=RPLP0 PE=1 SV=1 | 34 kDa | 96% |
| 78 | Cathepsin D OS=Homo sapiens OX=9606 GN=CTSD PE=1 SV=1 | 45 kDa | 96% |
| 79 | 40S ribosomal protein SA OS=Homo sapiens OX=9606 GN=RPSA PE=1 SV=4 | 33 kDa | 96% |
| 80 | Polyubiquitin-B OS=Homo sapiens OX=9606 GN=UBB PE=1 SV=1 | 26 kDa | 96% |
| 81 | 60 kDa heat shock protein, mitochondrial OS=Homo sapiens OX=9606 GN=HSPD1 PE=1 SV=2 | 61 kDa | 96% |
| 82 | Xaa-Pro dipeptidase OS=Homo sapiens OX=9606 GN=PEPD PE=1 SV=3 | 55 kDa | 96% |
| 83 | Plastin-2 OS=Homo sapiens OX=9606 GN=LCP1 PE=1 SV=6 | 70 kDa | 96% |
| 84 | Farnesyl pyrophosphate synthase OS=Homo sapiens OX=9606 GN=FDPS PE=1 SV=4 | 48 kDa | 96% |
| 85 | Adenylosuccinate synthetase isozyme 2 OS=Homo sapiens OX=9606 GN=ADSS2 PE=1 SV=3 | 50 kDa | 96% |
| 86 | Prolyl endopeptidase OS=Homo sapiens OX=9606 GN=PREP PE=1 SV=2 | 81 kDa | 96% |
| 87 | Reticulocalbin-1 OS=Homo sapiens OX=9606 GN=RCN1 PE=1 SV=1 | 39 kDa | 96% |
| 88 | Suprabasin OS=Homo sapiens OX=9606 GN=SBSN PE=1 SV=2 | 61 kDa | 96% |
| 89 | Serpin B12 OS=Homo sapiens OX=9606 GN=SERPINB12 PE=1 SV=1 | 46 kDa | 96% |
| 90 | MYG1 exonuclease OS=Homo sapiens OX=9606 GN=MYG1 PE=1 SV=3 | 42 kDa | 96% |
| 91 | Thyroid hormone receptor-associated protein 3 OS=Homo sapiens OX=9606 GN=THRAP3 PE=1 SV=2 | 109 kDa | 96% |
| 92 | Cartilage intermediate layer protein 1 OS=Homo sapiens OX=9606 GN=CILP PE=1 SV=4 | 133 kDa | 92% |
| 93 | Protein S100-A6 OS=Homo sapiens OX=9606 GN=S100A6 PE=1 SV=1 | 10 kDa | 86% |
| 94 | Vitamin K-dependent protein S OS=Homo sapiens OX=9606 GN=PROS1 PE=1 SV=1 | 75 kDa | 86% |
| 95 | Skin-specific protein 32 OS=Homo sapiens OX=9606 GN=XP32 PE=1 SV=1 | 26 kDa | 86% |
| 96 | P4K2A_HUMAN-DECOY | | 72% |
| 97 | TM141_HUMAN-DECOY | | 61% |
| 98 | CHD7_HUMAN-DECOY | | 59% |
| 99 | DX39A_HUMAN-DECOY | | 36% |
| 100 | NSRP1_HUMAN-DECOY | | 36% |
| 101 | Serotransferrin OS=Homo sapiens OX=9606 GN=TF PE=1 SV=3 | 77 kDa | 36% |
| 102 | Aldehyde dehydrogenase family 1 member A3 OS=Homo sapiens OX=9606 GN=ALDH1A3 PE=1 SV=2 | 56 kDa | 36% |
| 103 | Elongation factor Tu, mitochondrial OS=Homo sapiens OX=9606 GN=TUFM PE=1 SV=2 | 50 kDa | 36% |
| 104 | TP53-binding protein 1 OS=Homo sapiens OX=9606 GN=TP53BP1 PE=1 SV=2 | 214 kDa | 36% |
| 105 | Transcriptional adapter 2-beta OS=Homo sapiens OX=9606 GN=TADA2B PE=1 SV=2 | 48 kDa | 36% |
| 106 | Ecto-ADP-ribosyltransferase 4 OS=Homo sapiens OX=9606 GN=ART4 PE=2 SV=2 | 36 kDa | 36% |
| 107 | Hemicentin-1 OS=Homo sapiens OX=9606 GN=HMCN1 PE=1 SV=2 | 613 kDa | 36% |
| 108 | Voltage-dependent calcium channel gamma-6 subunit OS=Homo sapiens OX=9606 GN=CACNG6 PE=1 SV | 28 kDa | 36% |
| 109 | GRP1_HUMAN-DECOY | | 32% |
| 110 | Gasdermin-A OS=Homo sapiens OX=9606 GN=GSDMA PE=1 SV=4 | 49 kDa | 28% |
| 111 | Protocadherin gamma-B3 OS=Homo sapiens OX=9606 GN=PCDHGB3 PE=1 SV=4 | 101 kDa | 18% |
| 112 | Kininogen-1 OS=Homo sapiens OX=9606 GN=KNG1 PE=1 SV=2 | 72 kDa | 6% |
| 113 | Arfaptin-2 OS=Homo sapiens OX=9606 GN=ARFIP2 PE=1 SV=1 | 38 kDa | 6% |
| 114 | Protrudin OS=Homo sapiens OX=9606 GN=ZFYVE27 PE=1 SV=1 | 46 kDa | 6% |
| 115 | InaD-like protein OS=Homo sapiens OX=9606 GN=PATJ PE=1 SV=3 | 196 kDa | 6% |
| 116 | Poly(A) RNA polymerase, mitochondrial OS=Homo sapiens OX=9606 GN=MTPAP PE=1 SV=1 | 66 kDa | 6% |
| 117 | Keratin, type I cytoskeletal 28 OS=Homo sapiens OX=9606 GN=KRT28 PE=1 SV=2 | 51 kDa | 1% |
| 118 | Keratin, type I cytoskeletal 17 OS=Homo sapiens OX=9606 GN=KRT17 PE=1 SV=2 | 48 kDa | 1% |

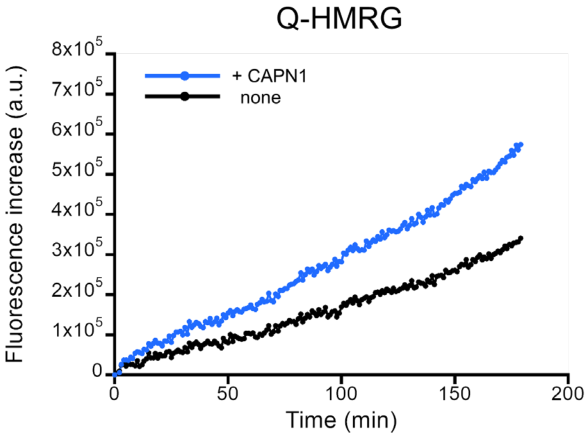


**Figure S5. Evaluation of the reactivity of Q-HMRG with CAPN1.** Time-dependent fluorescence changes of Q-HMRG in the presence or absence of CAPN1. The reaction of Q-HMRG with CAPN1 was performed in 25 mM Tris-HCl buffer (pH 7.5) containing 5 mM $CaCl_2$. [Q-HMRG] = 10 μM and [CAPN1] = 0.1 mg/mL. Fluorescence intensity was monitored using an EnVision plate reader (PerkinElmer). Filter settings were 485 (485/14 nm) for excitation and 535 (535/25 nm) for emission.

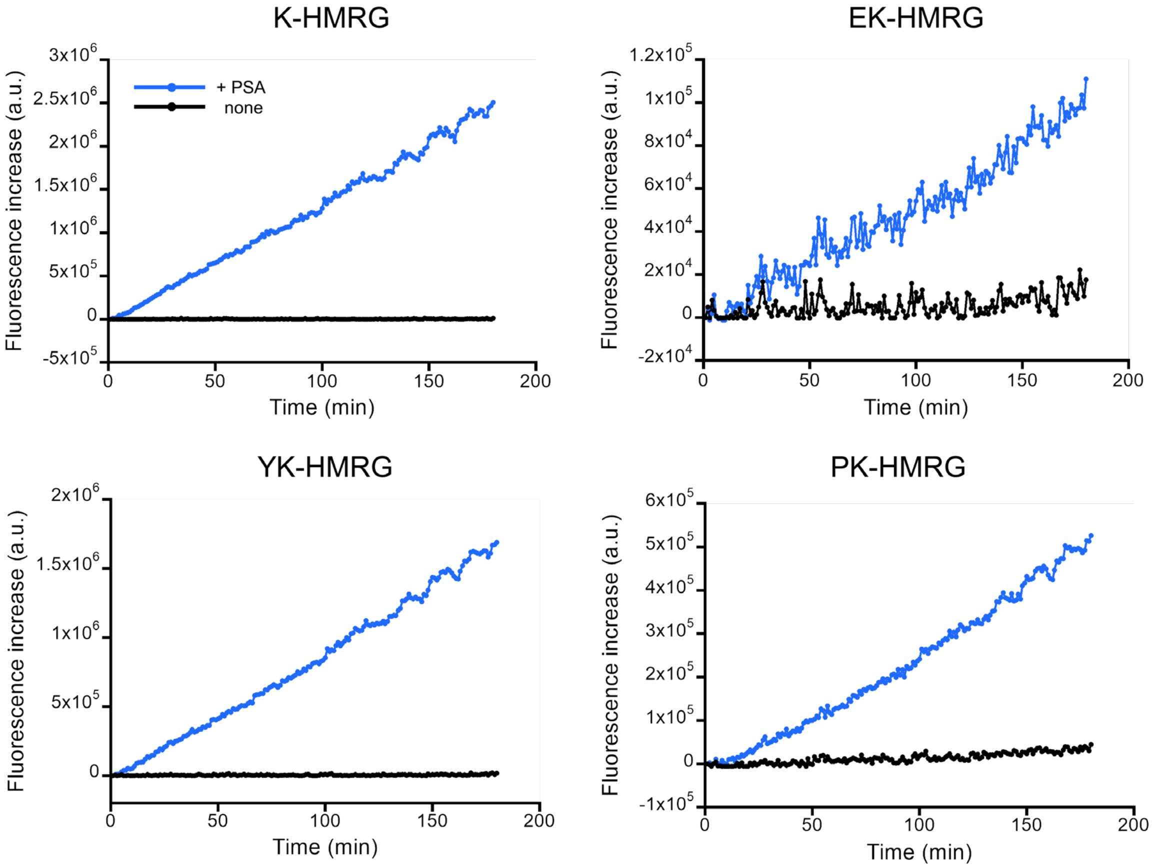


**Figure S6. Evaluation of the reactivity of fluorescence probes bearing lysine at the P1 position with PSA.** Time-dependent fluorescence changes of K-HMRG, EK-HMRG, YK-HMRG and PK-HMRG in the presence or absence of recombinant PSA. All four probes exhibited marked fluorescence increases in the presence of PSA, confirming their reactivity with PSA. All reactions were performed in PBS (−) containing 500 μM Triton X-100 at 37 °C for 120 min. [Fluorescence probe] = 1 μM and [PSA] = 250 ng/mL. Fluorescence intensity was monitored using an EnVision plate reader (PerkinElmer). Filter settings were 485 (485/14 nm) for excitation and 535 (535/25 nm) for emission.

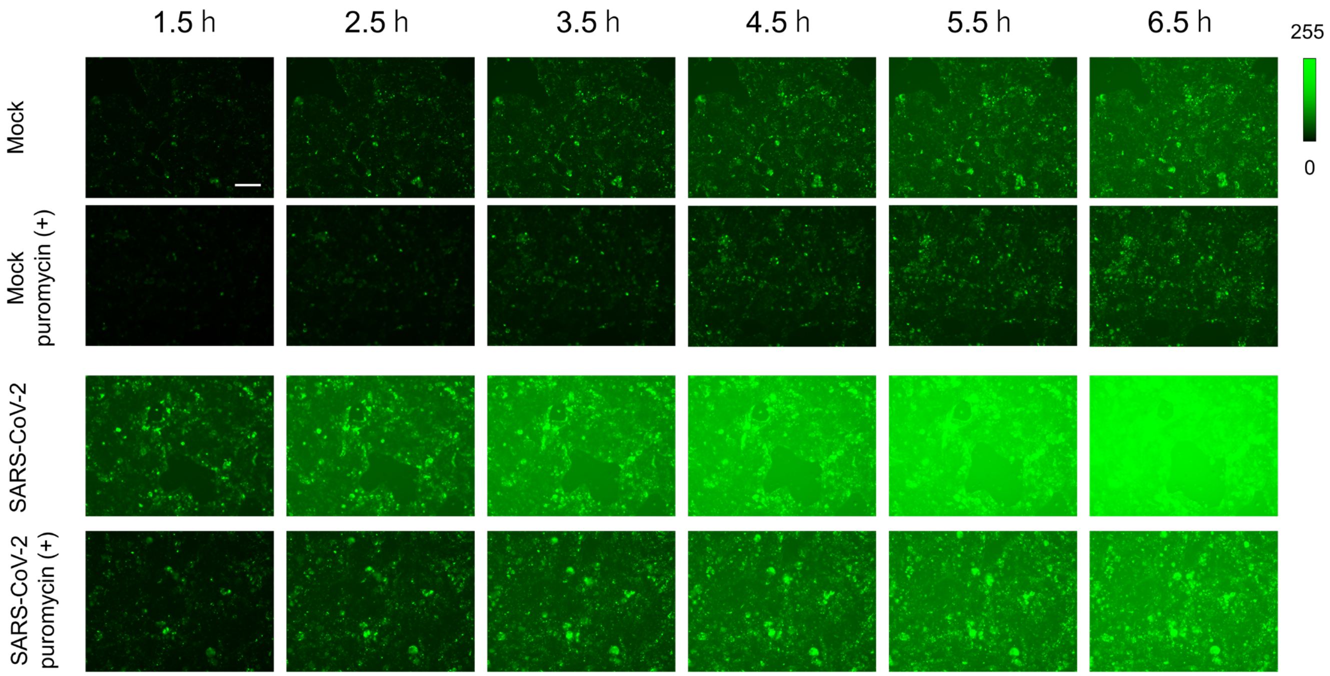


**Figure S7. Representative time-lapse fluorescence images of mock-infected and SARS-CoV-2-infected Calu-3 cells treated with YK-HMRG in the presence or absence of puromycin.** Images were acquired at 1.5-6.5 h after probe addition. Calu-3 cells were infected with SARS-CoV-2 at MOI of 10. [YK-HMRG] = 10 μM, [puromycin] = 50 μM. Scale bar, 200 μm.

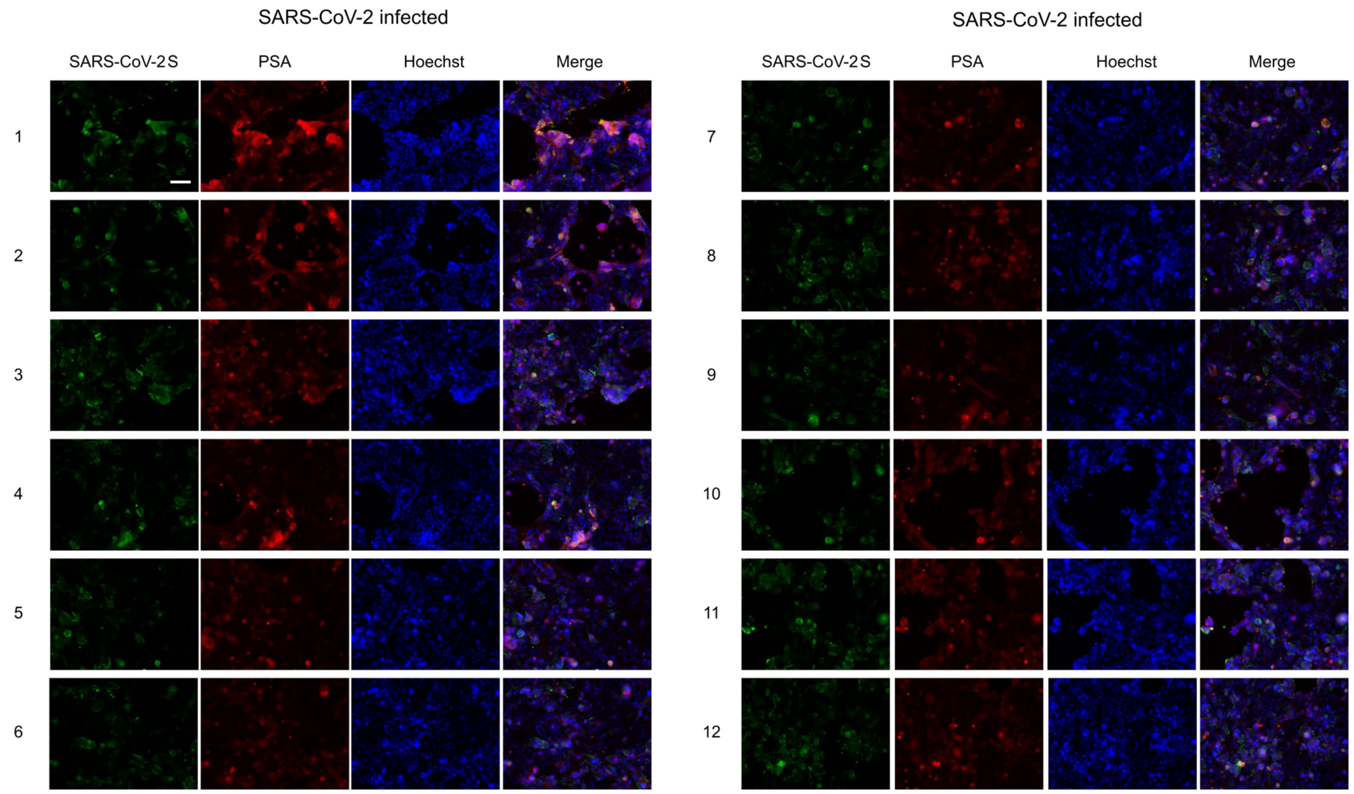


**Figure S8. Immunofluorescence analysis of PSA in SARS-CoV-2-infected Calu-3 cells.** Immunofluorescence staining of SARS-CoV-2 spike protein (green) and PSA (red) in SARS-CoV-2-infected Calu-3 cells. Calu-3 cells were infected with SARS-CoV-2 at MOI of 10. Nuclei were stained with Hoechst (blue). Merged images are shown in the rightmost panels. Scale bar, 100 μm.

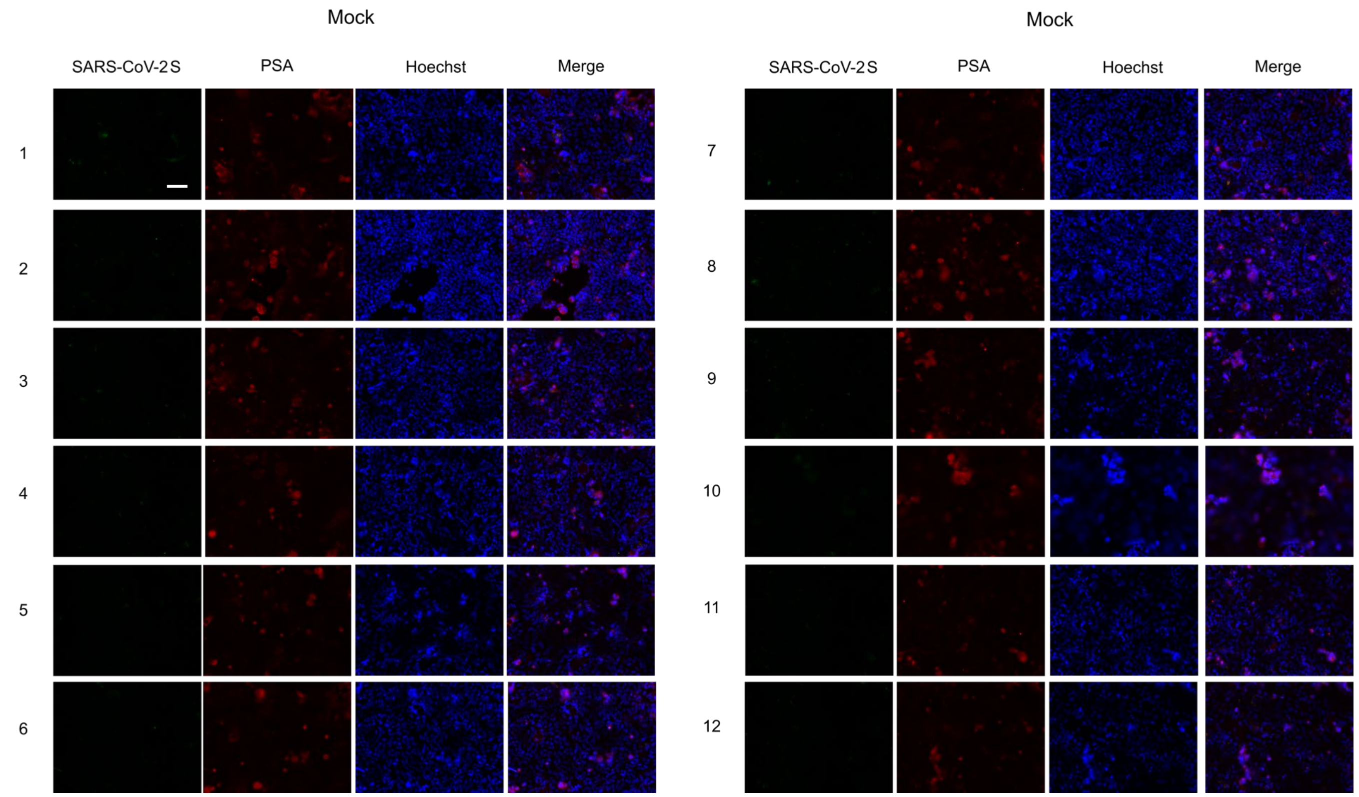


**Figure S9. Immunofluorescence analysis of PSA in mock-infected Calu-3 cells.** Immunofluorescence staining of SARS-CoV-2 spike protein (green) and PSA (red) in mock-infected Calu-3 cells. Nuclei were stained with Hoechst (blue). Merged images are shown in the rightmost panels. Scale bar, 100 µm.

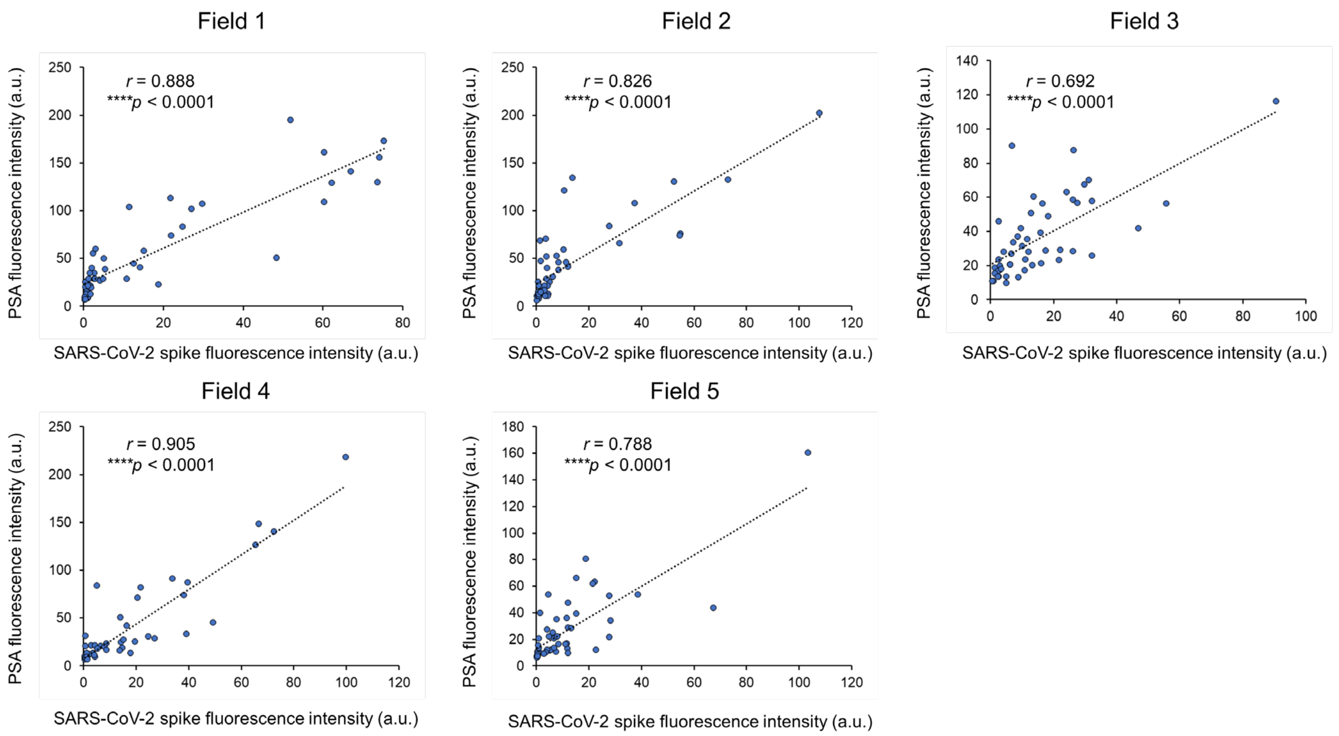


**Figure S10. Correlation between SARS-CoV-2 spike protein and PSA fluorescence intensities in individual imaging fields.** Scatter plots showing the correlations between SARS-CoV-2 spike protein and PSA fluorescence intensities in five imaging fields. Fifty cells were randomly selected from each field (n = 50 cells per field). Each dot represents an individual cell, and dotted lines indicate linear regression fits. Pearson's correlation coefficients ($r$) and corresponding $p$ values are indicated in each plot.

**Table S4. Translatomic and proteomic changes in CAPN1 and NPEPPS in SARS-CoV-2-infected Caco-2 cells.** Translatomic and proteomic data for CAPN1 and NPEPPS (encoding PSA) were extracted from Supplementary Tables 1 and 2, respectively, of Bojkova et al. (*Nature*, 2020). $Log_2$ ratios represent changes in protein synthesis rates (translatome) or total protein abundance (proteome) relative to mock-infected cells at the indicated time points after infection. P-values were calculated using two-sided, unpaired Student's t-tests (n = 3 independent biological samples). Statistical significance was assessed according to the criteria defined in the original study (*P < 0.05; ns, not significant).[1]

Translatome, SARS-CoV-2 vs Mock (Caco-2 cells)

| Protein | Time | $log_2$ ratio | P-value | Significant |
|---|---|---|---|---|
| CAPN1 | 2 h | 0.544 | 0.697 | ns |
| | 6 h | 0.176 | 0.894 | ns |
| | 10 h | −1.187 | 0.513 | ns |
| | 24 h | −0.767 | 0.63 | ns |
| NPEPPS (PSA) | 2 h | 1.26 | 0.388 | ns |
| | 6 h | −0.374 | 0.741 | ns |
| | 10 h | −0.219 | 0.858 | ns |
| | 24 h | 0.77 | 0.523 | ns |

Proteome, SARS-CoV-2 vs Mock (Caco-2 cells)

| Protein | Time | $log_2$ ratio | P-value | Significant |
|---|---|---|---|---|
| CAPN1 | 2 h | −0.0221 | 0.5411 | ns |
| | 6 h | −0.0159 | 0.0283 | * |
| | 10 h | 0.078 | 0.0317 | * |
| | 24 h | −0.0849 | 0.0523 | ns |
| NPEPPS (PSA) | 2 h | 0.03 | 0.585 | ns |
| | 6 h | 0.0316 | 0.2263 | ns |
| | 10 h | 0.0784 | 0.0937 | ns |
| | 24 h | −0.0061 | 0.845 | ns |

**Table S5. Transcriptomic and proteomic changes in CAPN1 and NPEPPS in SARS-CoV-2-infected A549-ACE2 cells.** Transcriptomic and proteomic data for CAPN1 and NPEPPS (encoding PSA) were extracted from Supplementary Tables 4 and 5, respectively, of Stukalov et al. (*Nature*, 2021). Log2 fold changes (log2FC) represent changes relative to mock-infected cells at the indicated time points after infection. Adjusted P-values are shown for transcriptomic data, whereas unadjusted P-values are shown for proteomic data. Statistical significance was assessed according to the criteria defined in the original study.[2] The "is_hit" column indicates whether each protein met the original study's criteria for differential regulation. ns, not significant.

Transcriptome, SARS-CoV-2 vs Mock (A549-ACE2 cells)

| Gene | Time | log2FC | Adjusted P-value | Significant |
|---|---|---|---|---|
| CAPN1 | 6 h | -0.0974 | 0.7195 | ns |
| | 12 h | -0.1795 | 0.2636 | ns |
| | 24 h | 0.0079 | 0.984 | ns |
| NPEPPS (PSA) | 6 h | 0.0772 | 0.8411 | ns |
| | 12 h | 0.1212 | 0.5501 | ns |
| | 24 h | -0.0033 | 0.9941 | ns |

Proteome, SARS-CoV-2 vs Mock (A549-ACE2 cells)

| Protein | Time | log2FC | P-value | is_hit |
|---|---|---|---|---|
| CAPN1 | 6 h | 0.064 | 0.06272 | FALSE |
| | 12 h | 0.0949 | 0.00311 | FALSE |
| | 24 h | 0.0888 | 0.01259 | FALSE |
| NPEPPS (PSA) | 6 h | 0.013 | 0.50555 | FALSE |
| | 12 h | 0.0123 | 0.71968 | FALSE |
| | 24 h | 0.0132 | 0.64782 | FALSE |

### References

(1) Bojkova, D.; Klann, K.; Koch, B.; Widera, M.; Krause, D.; Ciesek, S.; Cinatl, J.; Münch, C. Proteomics of SARS-CoV-2-infected host cells reveals therapy targets. *Nature* **2020**, *583* (7816), 469-472. DOI: 10.1038/s41586-020-2332-7.

(2) Stukalov, A.; Girault, V.; Grass, V.; Karayel, O.; Bergant, V.; Urban, C.; Haas, D. A.; Huang, Y.; Oubraham, L.; Wang, A.; et al. Multilevel proteomics reveals host perturbations by SARS-CoV-2 and SARS-CoV. *Nature* **2021**, *594* (7862), 246-252. DOI: 10.1038/s41586-021-03493-4.
